# Optogenetic manipulation of Gq- and Gi/o-coupled receptor signaling in neurons and heart muscle cells

**DOI:** 10.1101/2022.10.25.513732

**Authors:** Hanako Hagio, Wataru Koyama, Shiori Hosaka, Aysenur Deniz Song, Janchiv Narantsatsral, Koji Matsuda, Tomohiro Sugihara, Takashi Shimizu, Mitsumasa Koyanagi, Akihisa Terakita, Masahiko Hibi

**Affiliations:** Graduate School of Science, Nagoya University, Japan; Graduate School of Bioagricultural Sciences, Nagoya University; Institute for Advanced Research, Nagoya University; Graduate School of Science, Osaka Metropolitan University, Osaka, Japan

**Keywords:** Optogenetics, Bistable rhodopsin, G protein-coupled rhodopsin, G protein-coupled receptor, Locomotion, Cardiac contraction

## Abstract

G protein-coupled receptors (GPCRs) transmit extracellular signals into the cell depending on the type of G protein. To analyze the functions of GPCR signaling, we developed optogenetic tools using animal G protein-coupled bistable rhodopsins that can be controlled into active and inactive states by light irradiation. We expressed Gq- and Gi/o-coupled bistable rhodopsins in hindbrain reticulospinal V2a neurons, which are involved in locomotion, or in cardiomyocytes of zebrafish. Light stimulation of the reticulospinal V2a neurons expressing Gq-coupled spider Rh1 resulted in an increase in the level of cytoplasmic Ca^2+^ and evoked swimming behavior. Light stimulation of cardiomyocytes expressing the Gi/o-coupled mosquito Opn3, pufferfish TMT opsin, or lamprey parapinopsin induced cardiac arrest, and the effect was suppressed by treatment with pertussis toxin or barium, suggesting that Gi/o-dependent regulation of inward-rectifier K^+^ channels controls cardiac function. These data indicate that these rhodopsins are useful for optogenetic control of GPCR-mediated signaling in neurons and cardiomyocytes *in vivo*.

**Impact statement:** Animal G protein-coupled bistable rhodopsins can regulate Gq and Gi-mediated signaling in a light-dependent manner in neurons and cardiomyocytes, making them useful for analyzing roles of GPCR signaling *in vivo*.

## Introduction

G-protein coupled receptors (GPCRs) are responsible for transmitting extracellular signals into the cell. Many of them function as receptors for neurotransmitters or hormones, and activate coupled trimeric G proteins consisting of α, β, and γ subunits (*Hilger et al., 2018, Pierce et al., 2002, Rockman et al., 2002, Rosenbaum et al., 2009*). Upon activation of a GPCR, the α subunit (Gα) is converted from a GDP- to a GTP-bound form to regulate target proteins, while β and γ subunits are released from Gα as a complex (Gβγ) to control their own target proteins. GPCR regulates different signaling cascades depending on type of Gα that they bind (e.g., Gs, Gq, Gt, and Gi/o). Gs- and Gi/o-coupled GPCRs activate and inhibit, respectively cAMP-producing adenylyl cyclase (AC) via the Gα subunits. Gi/o-coupled GPCRs also regulate inward-rectifier K^+^ channels via the Gβγ subunit, increasing K^+^ efflux and thereby inducing hyperpolarization. On the other hand, Gq-coupled GPCRs, via their Gα subunits, activate phospholipase β (PLCβ) to generate inositol 1,4,5-triphosphate (IP3) and diacylglycerol (DAG) from phosphatidyl 4,5-bisphosphate (PIP2), subsequently elevating intracellular Ca^2+^ and activating protein kinase C (PKC). For example, in the central nervous system, the neurotransmitter glutamate binds to and activates GPCRs that are referred to as metabotropic receptors (GRMs), some of which function as Gq-coupled GPCRs (e.g., GRM1), and others as Gi/o-coupled GPCRs (e.g., GRM2, 3) (*Reiner and Levitz, 2018*). In the heart, noradrenaline binds to and activates the Gs-coupled β1 adrenergic receptor (β1AR), which increases myocardial contraction and heart rate (*de Lucia et al., 2018*), while acetylcholine binds to and activates the Gi/o-coupled muscarinic M2 receptor, which inhibits cardiac function (*Wess et al., 2007*). Although the functions of many GPCR signals have been studied, exactly in which cells, when, and how they function have not yet been fully elucidated. To solve these problems, it is necessary to precisely manipulate the location and timing of GPCR signaling.

Several techniques have been developed to control the activity and signaling of target cells. Chemogenetics using artificially designed GPCRs that are derived from muscarinic M3 receptor and can be activated by chemical ligands (Designer Receptor Exclusively Activated by Designer Drugs, DREADD) (*Armbruster et al., 2007, Kaganman, 2007, Roth, 2016, Wess et al., 2013*) has been used to control GPCR signaling, but the temporary and spatially precise control has been difficult. In contrast, optogenetics using rhodopsins, which bind to a chromophore retinal and can regulate their function in a light-sensitive manner, has been used to control and study cell functions. Light-gated microbial channelrhodopsins (e.g. ChR2) and light-driven microbial ion pump-type rhodopsins (e.g. halorhodopsin, NpHR) have been exploited to control the activities of neurons and/or cardiomyocytes (*Arrenberg et al., 2009, Arrenberg et al., 2010, Boyden et al., 2005, Deisseroth and Hegemann, 2017*). However, these rhodopsins induce depolarization or hyperpolarization of the membrane potential of cells in a light-stimulus-dependent manner at a precise timing and locations, but do not directly control GPCR signaling. On the other hand, animal rhodopsins are light-activated G protein-coupled proteins and can activate various signaling cascades like GPCRs for neurotransmitters and hormones, while displaying a diversity of wavelength sensitivity and G-protein selectivity (*Koyanagi et al., 2021, Koyanagi and Terakita, 2014, Terakita, 2005*).

Most animal rhodopsins bind to 11-*cis* retinal, which is isomerized to an all-*trans* form upon light absorption. This isomerization triggers a conformational change of rhodopsins and activates signal transduction cascades via the coupled G protein. Vertebrate visual rhodopsins release the chromophore all-*trans* retinal after light absorption and become an inactive form (bleach). The photoregeneration of these rhodopsins depends on the enzymes that generate 11-*cis* retinal, such as retinal isomerases, which are specifically expressed in photoreceptor organs (*Koyanagi et al., 2021, Koyanagi and Terakita, 2014, Terakita, 2005, Terakita et al., 2015*). Therefore, the photosensitivity of visual rhodopsins might not be very stable in cells other than photoreceptor organs. On the other hand, animal rhodopsins other than vertebrate visual rhodopsins retain 11-*cis* retinal and convert into photoproducts having the all-*trans* form (active state) upon light absorption, and these then revert to the original (inactive) dark state by subsequent light absorption, so they are bleach-resistant and are called bistable opsins (*Koyanagi et al., 2005, Koyanagi et al., 2021, Koyanagi and Terakita, 2014, Terakita and Nagata, 2014, Terakita et al., 2015, Tsukamoto and Terakita, 2010, Tsukamoto et al., 2005*). While chimeric optogenetic tools have been engineered using visual opsins with cytoplasmic loops and the C-terminal tail of adrenergic receptors (Opto-XRs) (*Airan et al., 2009, Kim et al., 2005, Siuda et al., 2015, Spangler and Bruchas, 2017*), bistable opsins have the advantage of stable optical control of GPCR signaling in various tissues.

A number of Gq-coupled bistable rhodopsin families have been identified as visual opsins in arthropods and molluscs, and as melanopsin in both vertebrates and invertebrates (*Koyanagi and Terakita, 2008*). Among them, jumping spider rhodopsin-1 (SpiRh1) was isolated from the jumping spider *Hasarius adansoni* and is reported to activate the Gq-signaling cascade in a green light-dependent manner (*Koyanagi et al., 2008, Nagata et al., 2012*). Mosquito Opn3 (MosOpn3) is an invertebrate homolog of vertebrate Opn3 (*Hill et al., 2002*). The Opn3 group contains multiple members including Opn3, originally called enecephalopsin, teleost multiple tissue (TMT) opsin, etc. (*Blackshaw and Snyder, 1999, Koyanagi et al., 2021, Moutsaki et al., 2003, Terakita, 2005, Terakita and Nagata, 2014*). When MosOpn3 was expressed in mammalian cultured cells, it bound to both 11-*cis* and 13-*cis* retinal (*Koyanagi et al., 2013*). MosOpn3 light-dependently activated Gi- and Go-type G proteins *in vitro* and initiated a Gi-signaling cascade in cultured cells (*Koyanagi et al., 2013*). Parapionopsin, which belongs to another group of bistable opsins, serves as a Gt-coupled opsin, like vertebrate visual opsins, and can also activate Gi-type G protein *in vitro* and in mammalian cultured cells (*Kawano-Yamashita et al., 2015, Koyanagi et al., 2021, Terakita et al., 2004, Tsukamoto et al., 2009*). The stable photoproduct (active form) of parapinopsin has its absorption maximum at ~500 nm, which is considerably distant from that of the dark state (~360 nm). Therefore, light illumination with different wavelengths was shown to switch on and off G protein-mediated signaling via parapinopsin *in vitro* and in cultured cells (*Kawano-Yamashita et al., 2015, Koyanagi et al., 2004, Wada et al., 2018*). Although MosOpn3 and lamprey parapinosin (LamPP) were used to suppress neuronal activities in a light stimulation-dependent manner (*Copits et al., 2021, Mahn et al., 2021*), it remains unclear whether they can control GPCR signaling in other types of cells and what mechanisms underlie optogenetic controls of GPCR signaling. In this study, we examined the optogenetic activity of animal bistable rhodopsins by expressing them in hindbrain reticulospinal V2a neurons that drive locomotion and cardiomyocytes in zebrafish.

## Results

### Activity of G protein-coupled bistable rhodopsins in human cells

To examine the activity of G protein-coupled rhodopsins in cells, we created two DNA constructs that expressed a rhodopsin and a fluorescent protein as a fusion protein, or that expressed a carboxy-terminal epitope-tagged rhodopsin and a fluorescent protein separately using a viral 2A (P2A) peptide system. We first expressed a fusion protein of Gq-coupled SpiRh1 (*Koyanagi et al., 2008, Nagata et al., 2012*) and TagCFP (SpiRh1-TagCFP), or Flag-tagged SpiRh1 and TagCFP separately (SpiRh1-P2A-TagCFP), in human embryonic kidney (HEK293S) cells (Figure 1A). Light stimulation increased intracellular Ca^2+^ at a much higher level for SpiRh1-P2A-TagCFP-expressing cells than SpiRh1-TagCFP-expressing cells, suggesting that the expression level and/or activity of bistable rhodopsins is higher with a small peptide (epitope)-tagged protein than with a large fluorescent-fused protein. The light stimulation-dependent increase in intracellular Ca^2+^ with SpiRh1 was suppressed by treatment with a Gαq inhibitor YM254890 (Figure 1D), confirming that SpiRh1 mediates Gq-mediated signaling. We used three different epitope tags (influenza virus hemagglutinin [HA], Myc [MT], and Flag tags) to express bistable rhodopsins, finding no significant differences between them (data not shown). We thus chose to use a Flag tag hereafter. We created similar expression constructs for bistable Gq- and Gi/o-coupled rhodopsins from various invertebrate and vertebrate animals listed in Table 1 and expressed these rhodopsins in HEK293S cells. These included Gq-coupled SpiRh1[S186F], a SpRh1 mutant that has a maximal sensitivity in the UV region (*Nagata et al., 2019*) as well as Gi/o-coupled MosOpn3 (carboxy-terminal truncated MosOpn3 was used) (*Koyanagi et al., 2013*), pufferfish TMT opsin (PufTMT) (*Koyanagi et al., 2013*), lamprey parapinopsin (LamPP) (*Koyanagi et al., 2004*), and zebrafish parapinopsin1 (ZPP1) (*Koyanagi et al., 2015*). Stimulation of SpiRh1 and SpiRh1[S186F]-expressing cells with 500 and 410 nm light, respectively, increased intracellular Ca^2+^ (Figure 1B). Light stimulation of cells expressing MosOpn3, PufTMT, LamPP, or ZPP1 with 500 (for MosOpn3 and PufTMT) or 410 nm (for LamPP and ZPP1) light reduced intracellular cAMP levels to similar extents (Figure 1C). These data indicate that these Flag-tagged G protein-coupled rhodopsins can be used for optogenetic manipulation of Gq- and Gi/o-mediated signaling in vertebrate cells.

**Figure 1.**
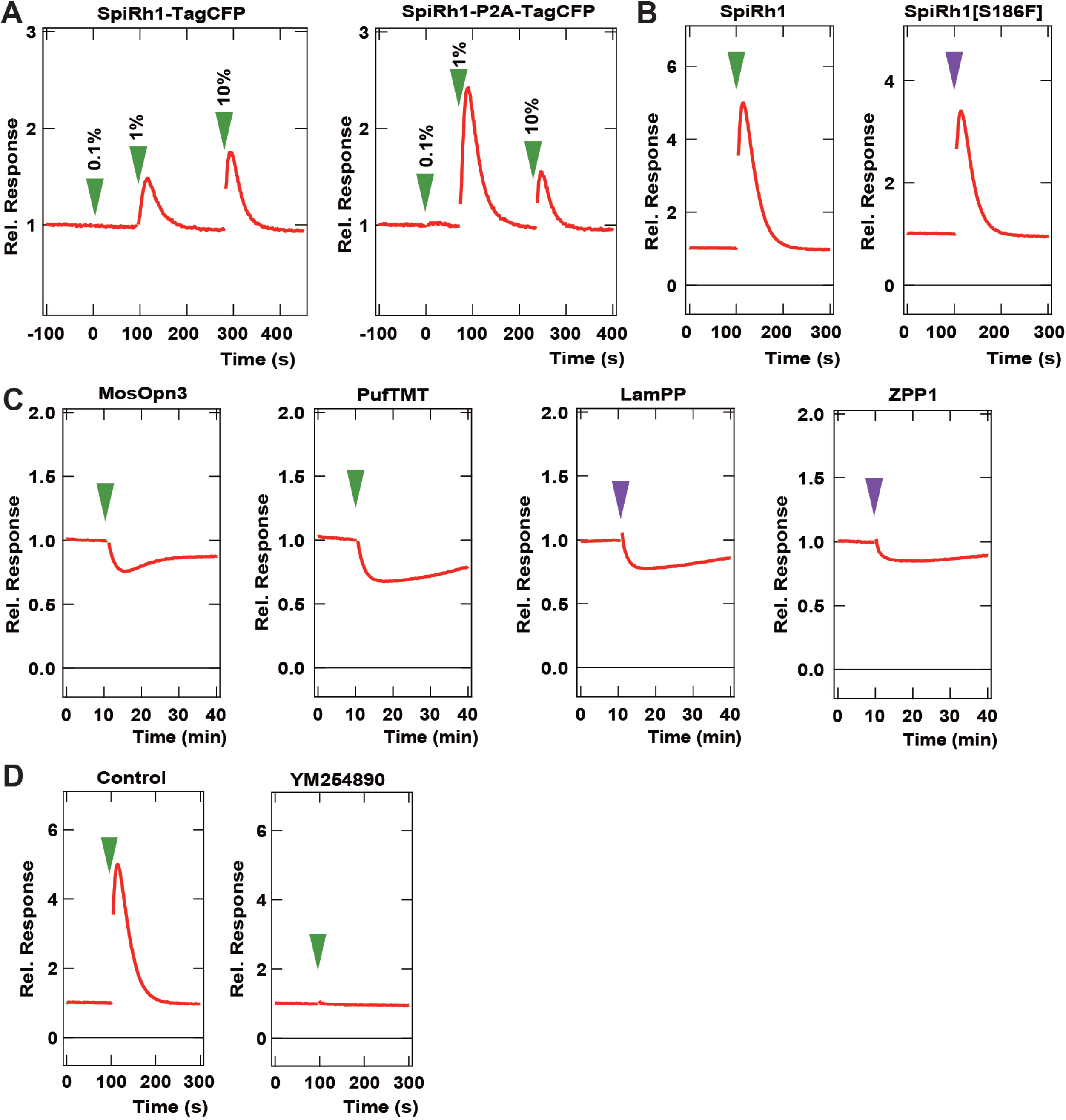
Activity of G-coupled bistable rhodopsins in HEK293S cells. (A) Comparison of optogenetic activities of Gq-coupled Spider Rh1 (SpiRh1) expressed using TagCFP fusion protein and the P2A-TagCFP system. HEK293S cells were transfected with an expression plasmid for the fusion protein of SpiRh1 and TagCFP (SpiRh1-TagCFP, left panel), or for that of Flag-tagged SpiRh1, porcine teschovirus 2A peptide, and TagCFP (Spi-P2A-TagCFP, right panel). Transfected cells were incubated with 11-*cis* retinal and stimulated by different intensities of 500 nm light (0.1%, 1%, or 10% of the light intensity, with 0.106 mW/mm^2^ as 100%). Intracellular Ca^2+^ concentration was measured by using aequorin m2 and is indicated as a ratio to the unstimulated state in the graphs. (B) Comparison of activities of Flag-tagged SpiRh1 and SpiRh1 [S186F]. Transfected cells were stimulated by 500 nm (green arrow, 0.106 mW/mm^2^) or 410 nm (purple arrow, 0.0194 mW/mm^2^) light and intracellular Ca^2+^ concentration was measured. (C) Light-stimulus-dependent reduction of intracellular cAMP level by Gi/o-coupled Mosquito Opn3 (MosOpn3), Pufferfish TMT (PufTMT), Lamprey PP (LamPP), and zebrafish PP1 (ZPP1). HEK293S cells were transfected with expression plasmids for flagged-tagged Gi/o rhodopsins. Transfected cells were incubated with 11-*cis* retinal and stimulated by 500 nm (green arrow) or 410 nm (purple arrow) light. Intracellular cAMP concentration was measured with GloSensor 20F and is indicated as a ratio to the unstimulated state. (D) Effects of Gαq inhibitor YM254890 on SpiRh1. HEK293S transfected with an expression plasmid for SpiRh1 were incubated with 11-*cis* retinal alone (left panels) or with 11-*cis* retinal and YM254890 (right panels), and were stimulated by 500 nm light. Intracellular Ca^2+^ concentration was measured.

**Table 1.**
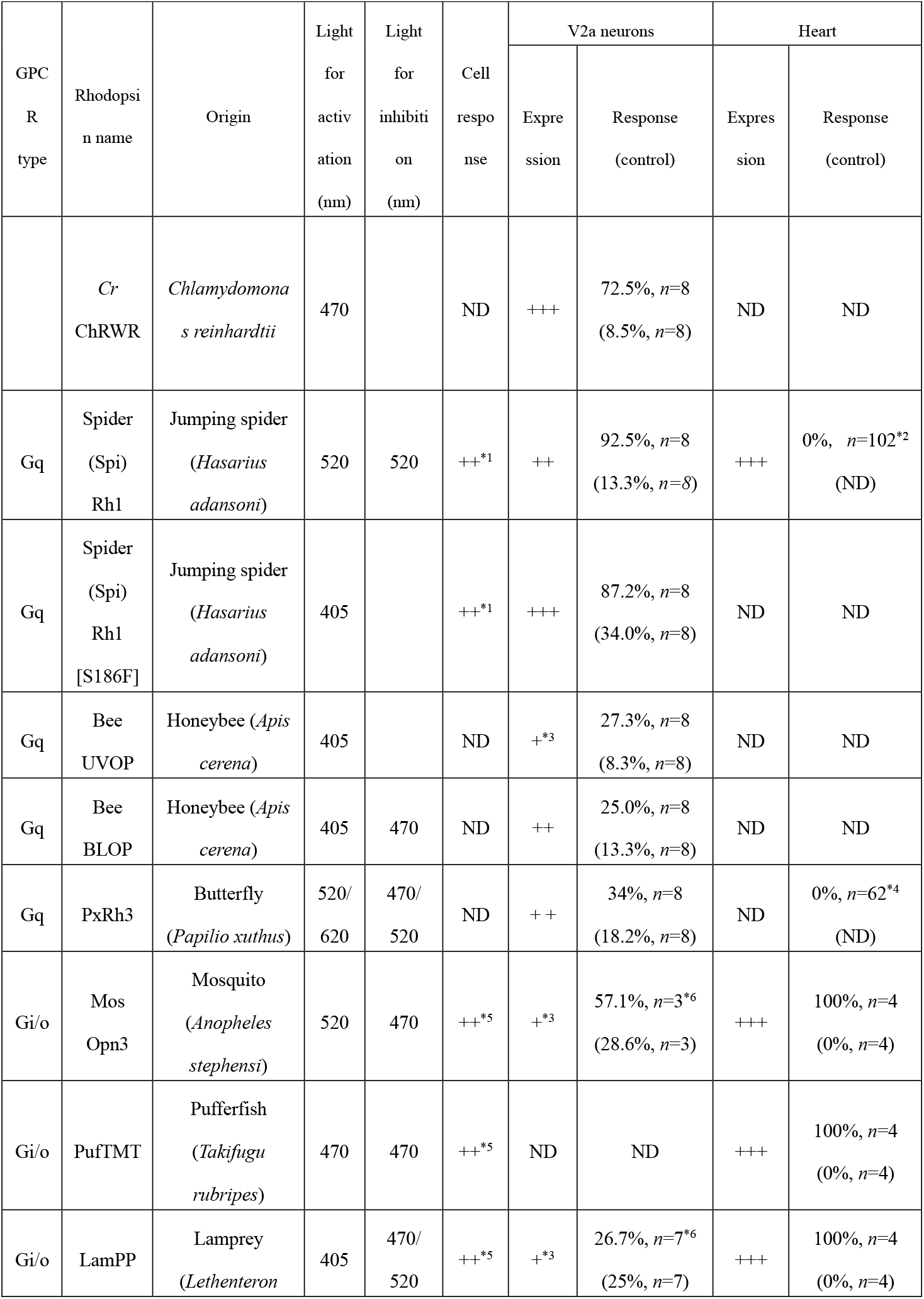
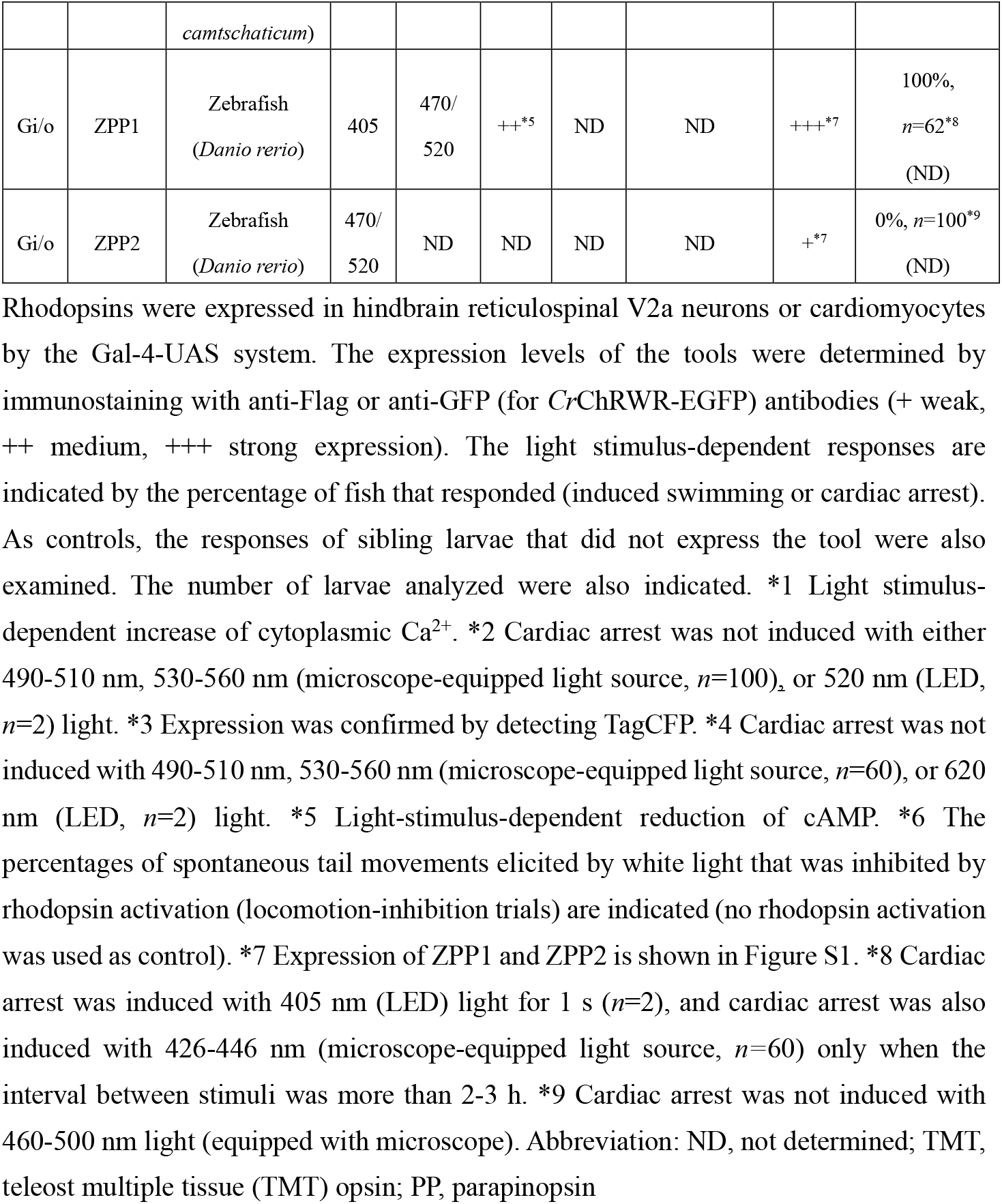
Summary of optogenetic tools

### Optogenetic activation of zebrafish locomotion circuit by Spider Rh1

To evaluate the optogenetic activities of the G protein-coupled rhodopsins *in vivo*, we expressed these rhodopsins in either hindbrain reticulospinal V2a neurons that were reported to drive locomotion (*Kimura et al., 2013*) or in cardiomyocytes of zebrafish larvae by using the Gal4-UAS system. Transgenic zebrafish *Tg(vsx2:GAL4FF)*, which is also known as *Tg(chx10:GAL4)*, express a modified version of the transcriptional activator GAL4-VP16 (*Asakawa et al., 2008*) in hindbrain reticulospinal V2a neurons (*Kimura et al., 2013*). We generated stable transgenic lines *Tg(UAS-hsp70l:opto-tool)* that can express Flag-tagged rhodopsins with P2A-TagCFP under the control of 5xUAS (upstream activating sequences of the yeast *Gal1* gene) and the zebrafish *hsp70l* promoter (*Muto et al., 2017*), and mCherry in the heart. We used an EGFP fusion protein of the channelrhodopsin wide receiver (ChRWR), which is a chimeric protein of *Chlamydomonas reinhardtii* channelrhodopsins ChR1 and ChR2, as a positive control (*Kimura et al., 2013, Umeda et al., 2013, Wang et al., 2009*). We crossed *Tg(vsx2:GAL4FF);Tg(UAS:RFP)* and *Tg(UAS-hsp70l:opto-tool)* to express various G protein-coupled rhodopsins and ChRWR-EGFP listed in Table 1. The expression of these rhodopsins was examined by TagCFP or fused EGFP expression in 3-days post fertilization (3-dpf) Tg larvae, and was further analyzed by immunohistochemistry (Figure 2B, Table 1). Since transgene-mediated protein expression depends on the nature of the introduced gene, the transgene-integrated sites and copy number, we established multiple Tg lines and analyzed stable Tg lines (F1 or later generations) with the highest tool expression and relatively close expression levels. We irradiated a hindbrain area of 3-dpf Tg larvae expressing the G protein-coupled rhodopsins with light of wavelength near their absorption maxima to stimulate each rhodopsin, using a patterned illuminator (Figure 2A, Table 1). Tail movements after light stimuli were monitored (Figure 2C-E, Movie 1-3). The rate at which light irradiation was able to induce tail movements (locomotion rate, Figure 3A), the time from irradiation to the onset of tail movements (latency, Figure 3B), the duration of tail movements (Figure 3C), and the amplitude of tail movements, were measured (Figure 3D).

**Figure 2.**
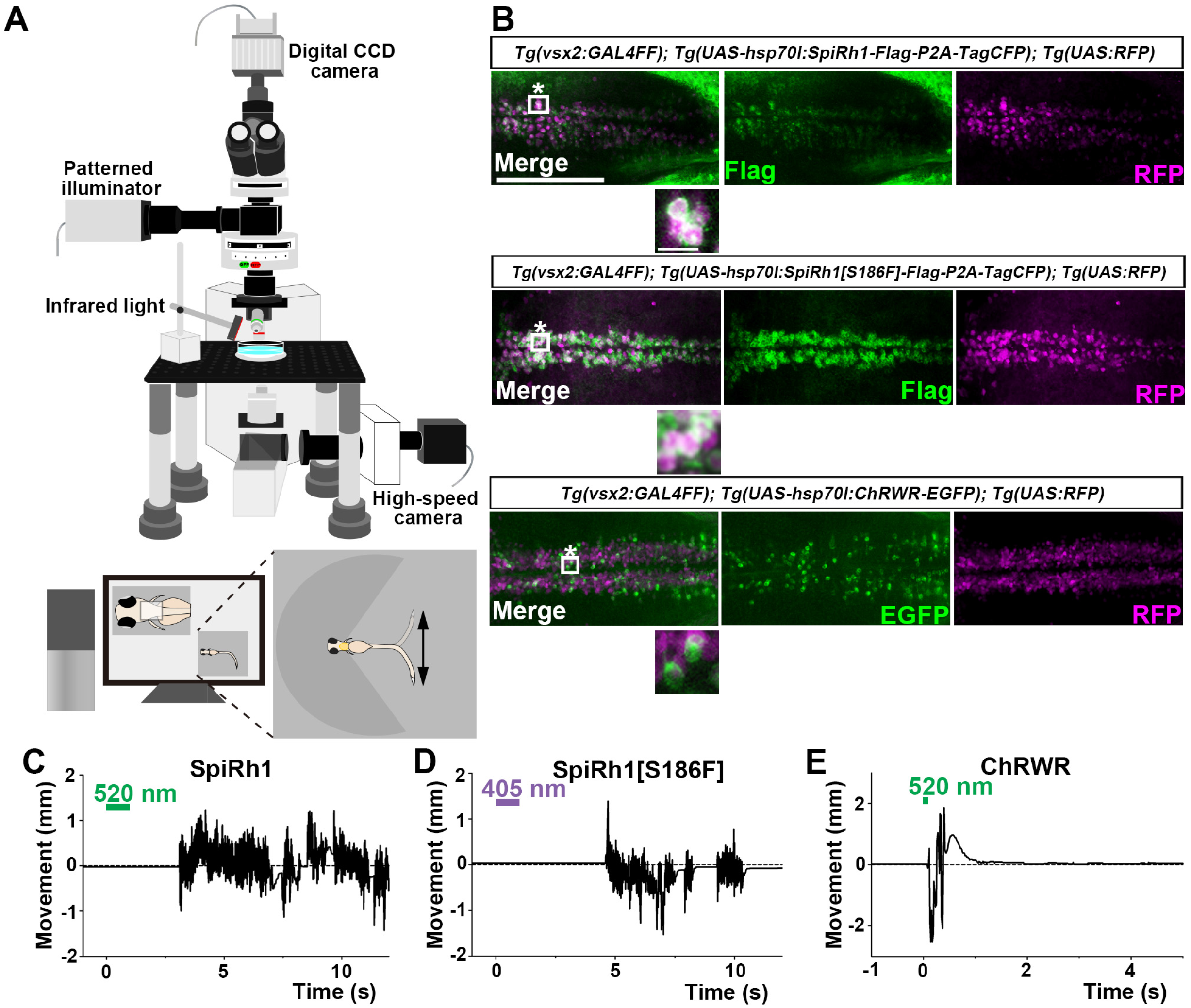
Activation of reticulospinal V2a neurons by Gq-coupled bistable rhodopsins. (A) Schematic of experimental devices for induction of swimming behavior and a larva embedded in agarose. The hindbrain region was irradiated with light by using a patterned illuminator. Tail (caudal fin) movements were monitored by a high-speed camera with infrared light. (B) Expression of SpiRh1, SpiRh1[S186F], and channel rhodopsin wide receiver (ChRWR) in hindbrain reticulospinal V2a neurons. 3-dpf (days post fertilization) *Tg(vsx2:GAL4FF);Tg(UAS-hsp70l:SpiRh1-Flag-P2A-TagCFP, myl7:mCherry);Tg(UAS:RFP), Tg(vsx2:GAL4FF);Tg(UAS-hsp70l:SpiRh1[S186F]-Flag-P2A-TagCFP, myl7:mCherry);Tg(UAS:RFP)* and *Tg(vsx2:GAL4FF);Tg(UAS:ChRWR-EGFP);Tg(UAS:RFP)* larvae were fixed and stained with anti-Flag or anti-GFP (EGFP, green), and anti-DsRed (RFP, magenta) antibodies. Inset: higher magnification views of the boxed areas showing double-labeled neurons. In the inset, fluorescence signal intensities were modified to compare the subcellular localization of the tools. (C, D, E) Tail movements of 3-dpf Tg larvae expressing SpiRh1 (C), SpiRh1 [S186F] (D), and ChRWR (E) in the reticulospinal V2a neurons after light stimulation. The hindbrain area was stimulated with light (0.4 mW/mm^2^) of wavelength of 520 nm (for SpiRh1), 405 nm (for SpiRh1[S186F]), and 470 nm (for ChRWR) for 1 s (for SpiRh1 and SpiRh1[S186F]) or 100 ms (for ChRWR). Typical movies are shown in Movie 1-3. Scale bar = 150 μm in (B), 10 μm in in the insets of (B).

**Figure 3.**
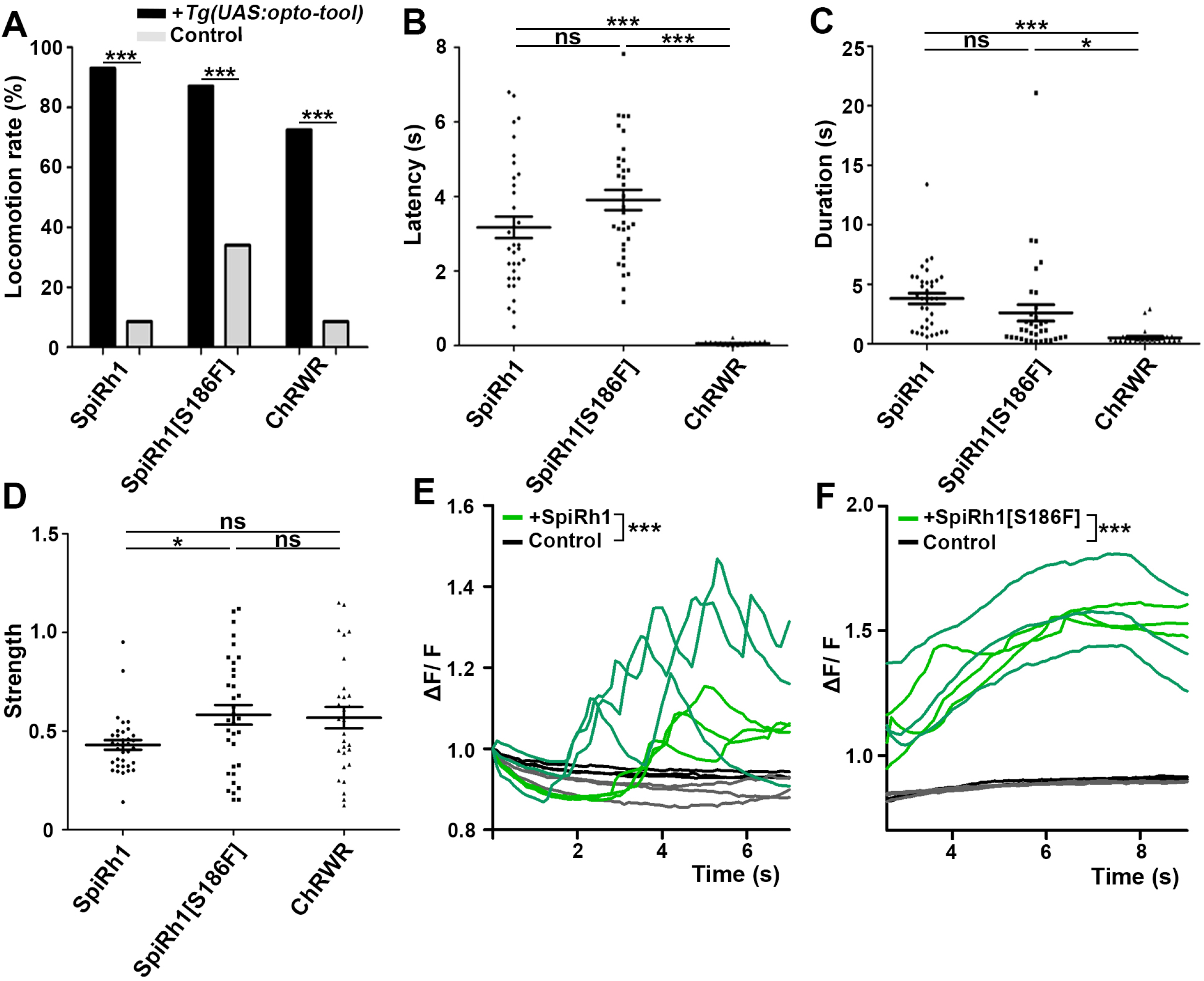
Locomotion induced by SpiRh1, SpiRh1[S186F], and ChRWR. (A) Light stimulus-dependent locomotion rates of 3-dpf Tg larvae expressing SpiRh1, SpiRh1[S186F] and ChRWR in hindbrain reticulospinal V2a neurons. Sibling larvae that did not express the tools were used as controls. Six consecutive stimulus trials were analyzed for eight larvae of each Tg line. The percentage of trials in which tail movement was elicited was defined as locomotion rate. The locomotion rates for SpiRh1, SpiRh1[S186S], and ChRWR were 92.5% (control 13.3%), 87.2% (control 34.0%), and 72.5% (control 8.5%), respectively. *** *p* < 0.001, Fisher’s exact test. (B, C, D) Light stimulus-evoked tail movements of latency (B), duration (C), and strength (D). The time from the start of light irradiation to the first tail movement was defined as latency (s), and the time from the beginning to the end of the first tail movement was defined as duration (s). The maximum distance the caudal fin moved from the midline divided by the body length was measured as strength. The latency of SpiRh1, SpiRh1[S186S], and ChRWR was 3.17±0.289, 3.91±0.271, and 0.0617±0.00806 s, respectively. The duration of SpiRh1, SpiRh1[S186S], and ChRWR was 3.80±0.449, 2.59±0.692, and 0.504±0.122 s, respectively. The strength of SpiRh1, SpiRh1[S186S], and ChRWR was 0.430±0.0246, 0.582±0.0498, and 0.568±0.0541 mm, respectively. (E, F) Light-evoked Ca^2+^ increased with SpiRh1 (E) and SpiRh1[S186F] (F) in hindbrain V2a neurons. 3-dpf *Tg(vsx2:GAL4FF);Tg(UAS-hsp70l:SpiRh1-Flag-P2A-TagCFP, myl7:mCherry);Tg(UAS-hsp70l:GCaMP6s)* and *Tg(vsx2:GAL4FF);Tg(UAS-hsp70l:SpiRh1[S186F]-Flag-P2A-TagCFP, myl7:mCherry);Tg(UAS-hsp70l:GCaMP6s)* larvae were used. Sibling larvae that expressed GCaMP6s but did not express SpiRh1 or SpiRh1[S186F] were used as controls. The hindbrain area was irradiated and GCaMP6s fluorescence was detected with a fluorescence detection filter (excitation 470-495 nm, emission 510-550 nm) for SpiRh1. For SpiRh1[S186F], GCaMP6s fluorescence was detected after irradiation with 405 nm light for 1 s. Two larvae for each condition (SpiRh1, SpiRh1[S186F], and controls) were analyzed and three consecutive trials were analyzed. The change in fluorescence intensity of GCaMP6s (ΔF/F) is indicated as a ratio to the fluorescence intensity at the start of stimulation (E) for SpiRh1 and before (F) the start of stimulation with 405 nm light for SpiRh1[S186F]. The ΔF/F of Tg larvae expressing SpiRh1 or SpiRh1[S186F] is indicated by green lines whereas that of control larvae is indicated by black lines. Ca^2+^ responses were significantly higher in Tg larvae expressing SpiRh1 and SpiRh1[S186F] than control larvae. *** *p* < 0.001, linear mixed-effects model.

Among the G protein-coupled rhodopsins examined, we found that SpiRh1 and SpiRh1[S186F] were most potent in inducing tail movements. Immunohistochemistry with anti-Flag or GFP antibodies revealed ChRWR expression on the cell surface of hindbrain reticulospinal V2a neurons was mosaic due to methylation-dependent silencing of the UAS system (*Akitake et al., 2011*), while SpiRh1 and SpiRh1[S186F] were uniformly expressed in these neurons, with relatively low SpiRh1 expression compared to SpiRh1[S186F] (Figure 2B). As was previously reported (*Kimura et al., 2013*), light stimulation of reticulospinal V2a neurons with ChRWR for 100 ms immediately evoked tail movements (Figure 2E). Activation with SpiRh1 and SpiRh1[S186F] required longer stimulation (1 s) and 3-5 s to initiate tail movements (Figure 2C, D, E, Figure 3B, Movie 1-3). However, stimulation with SpiRh1 and SpiRh1[S186F] elicited tail movements for a longer period than ChRWR (Figure 3C). Light stimulation of control sibling larvae that did not express the rhodopsins scarcely induced tail movements, although stimulation with 405 nm light induced locomotion at a low frequency. This is possibly due to toxicity of low-wavelength light, close to the UV range. The data suggests that optical activation of reticulospinal V2 neurons with SpiRh1 and SpiRh1[S186F] is robust and long-lasting, although it requires longer stimulation and longer latency than channelrhodopsin. In G protein-mediated signaling, it is generally accepted that Gq activates PLCβ and thereby generates IP3, which induces Ca^2+^ influx from the endoplasmic reticulum. To examine the level of intracellular Ca^2+^ level, we expressed SpiRh1 or SpiRh1[S186F] with GCaMP6s in hindbrain reticulospinal V2a neurons. We found that light stimulation with these Gq-coupled rhodopsins increased the intracellular Ca^2+^ level in these neurons (Figure 3E, F, Movie 4, 5).

### Optogenetic manipulation of zebrafish heart by Gi/o-coupled rhodopsins

Gi/o-coupled bistable rhodopsins MosOpn3 and LamPP were used to suppress neuronal activities (*Copits et al., 2021, Mahn et al., 2021*). We next examined whether they could be used to control cardiomyocyte function *in vivo*. We expressed the Gi/o-coupled rhodopsins by using *Tg(myl7:GAL4FF)*, in which GAL4FF was expressed under the control of the promoter of the cardiac myosin light chain gene *myl7*. By crossing *Tg(UAS-hsp70l:opto-tool)*, we expressed Gi/o-coupled rhodopsins listed in Table 1. We again established multiple Tg lines and analyzed stable Tg lines with the highest tool expression and relatively close expression levels. Immunohistochemical staining revealed comparable expression of these Gi/o-coupled rhodopsins in zebrafish cardiomyocytes (Figure 4A, S1, Table 1). We irradiated the entire heart area of 4-dpf Tg fish expressing the Gi/o-coupled rhodopsins with light of appropriate wavelengths for 1 s. Videos of heartbeats (HBs) before and after light stimulation were recorded (Movie 6 for MosOpn3, Movie 7 for PufTMT, Movie 8 for LamPP). HBs were analyzed and heart rates were calculated (Figure 4B, C). Activation of the Gi/o-coupled rhodopsins MosOpn3, PufTMT, and LamPP in the heart led to cardiac arrest within 1 s in all Tg larvae examined, but not in control sibling larvae (Figure 4 B, C, D, E, Movie 6-8). The first HB occurred about 10 s after cardiac arrest, but it took at least 1 min (sometimes a few minutes) to return to normal HBs (Figure 4B, C, D, E, F, Movie 6-8). Optical activation of ZPP1 in the heart induced cardiac arrest for several seconds, while light stimulus-dependent cardiac arrest was not observed unless the time interval between stimuli exceeded 2-3 h (Table 1). The data suggest that MosOpn3, PufTMT, and LamPP are efficient optogenetic tools to control the function of cardiomyocytes in zebrafish.

**Figure 4.**
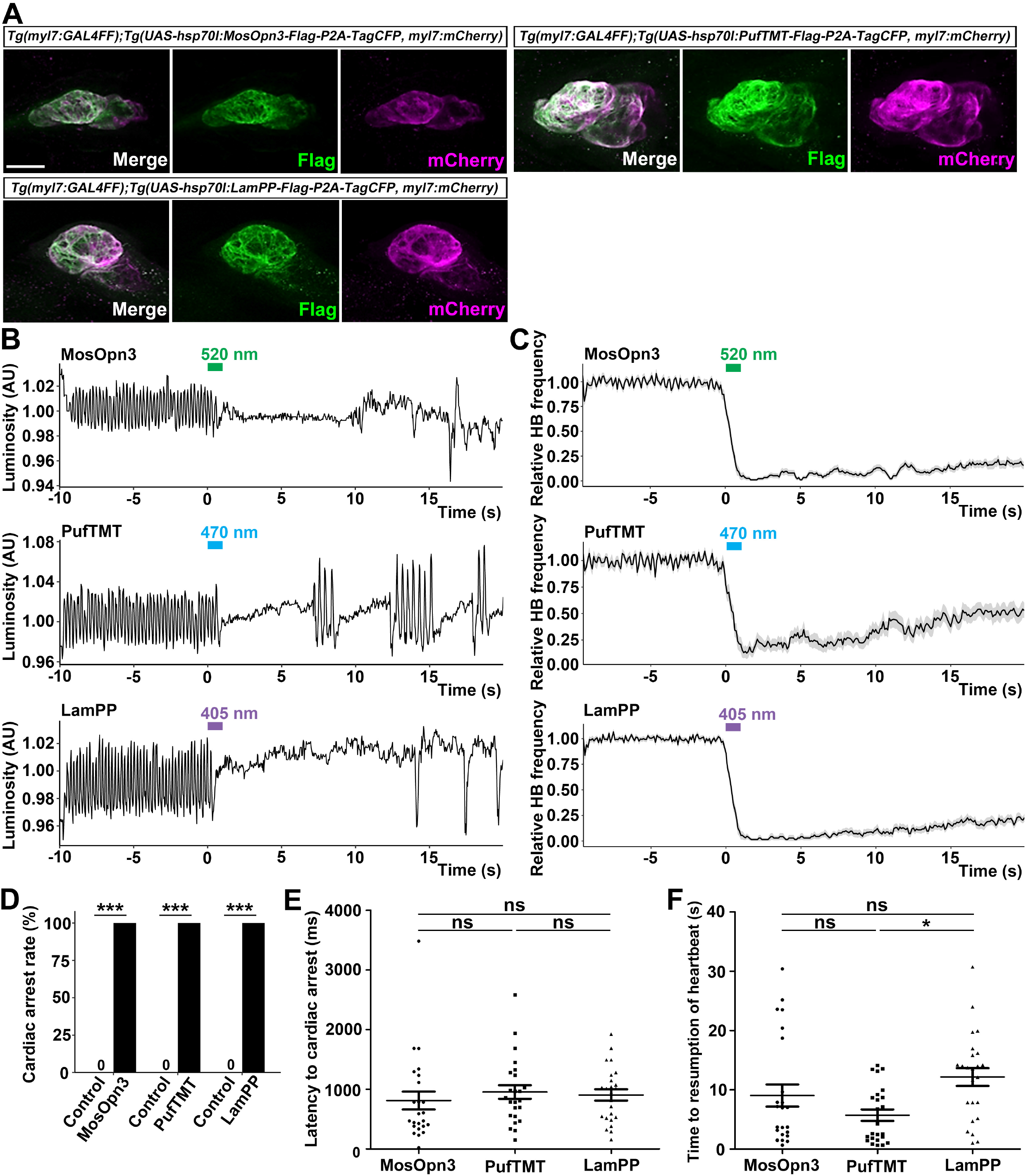
Inhibition of cardiomyocytes by Gi/o-coupled bistable rhodopsins. (A) Expression of Gi/o-coupled rhodopsins MosOpn3, PufTMT, and LamPP in zebrafish cardiomyocytes. 4-dpf *Tg(myl7:GAL4FF);Tg(UAS:opto-tool-Flag-P2A-TagCFP, myl7:mCherry)* larvae were fixed and stained with anti-Flag (green) and anti-DsRed (mCherry: magenta). (B, C) Heartbeat (HB) monitoring by change in luminosity (AU: arbitrary units) (B) and the average relative HB frequency (C) of four larvae expressing MosOpn3, PufTMT, and LamPP in cardiomyocytes. The heart area of larvae expressing MosOpn3, PufTMT, and LamPP in cardiomyocytes were stimulated with 520, 470, and 405 nm light (0.5 mW/mm^2^), respectively, for 1 s. Six consecutive stimulus trials were analyzed for four rhodopsin-expressing larvae of each Tg line. Typical HB data are shown in (B) and the average HB frequency for 24 trials are shown in (C). (D) Cardiac arrest rates. Six consecutive stimulus trials were analyzed for four rhodopsin-expressing larvae and four control larvae of each Tg line (MosOpn3, PufTMT, and LamPP). The cardiac arrest rates for MosOpn3, PufTMT, and LamPP were all 100% (control 0%). *** *p* < 0.001, Fisher’s exact test. (E, F) Latency to cardiac arrest (E), and time to resumption of HBs (F) with MosOpn3, PufTMT, and LamPP by light stimulation. Latencies to cardiac arrest in MosOpn3, PufTMT, and LamPP were 812±147, 955±112, and 905±93 ms, respectively. Times to resumption of HB in MosOpn3, PufTMT, and LamPP were 8.8±1.9, 5.7±1.0, and 12.1±1.4 s, respectively. * *p* < 0.05, One-way ANOVA with Tukey’s post hoc test. Scale bar = 50 μm in (A).

Bistable rhodopsins convert to active states upon light stimulation, and then revert to the original inactive dark state by subsequent light absorption. Thus, the activity of these rhodopsins can be switched off by light stimulation after activation. The activation and inactivation wavelengths are close to each other for MosOpn3 and PufTMT, but apart for LamPP (Table 1). We assessed inactivation of the Gi/o-coupled rhodopsins by sustained light stimulation. We expressed MosOpn3, PufTMT, or LamPP together with GCaMP6s in cardiomyocytes, and simultaneously monitored intracellular Ca^2+^ and HBs. Continuous stimulation of MosOpn3 with 470 nm light initially led to cardiac arrest and a reduction in intracellular Ca^2+^ concentration in both the atrium and ventricle of the heart within 20 s. However, HBs resumed and intracellular Ca^2+^ gradually increased around 40 s during light stimulation, and the HBs returned to a steady state at around 70 s (Figure 5A, Movie 9). Continuous light stimulation of PufTMT in the heart caused cardiac arrest and a reduction in intracellular Ca^2+^ concentration in about 5 s, followed by resumption of HBs in 5-10 s, and the return to a steady state at around 20 s (Figure 5B). These data suggest that sustained light stimulation can activate and subsequently inactivate MosOpn3 and PufTMT due to light adaptation. Stimulation of LamPP with 405 nm light in the heart led to cardiac arrest and a reduction in Ca^2+^, while subsequent sustained stimulation with 470 nm light recovered both heart rate and Ca^2+^ concentration (Figure 5C, D, E, Movie 10). Therefore, the activity of LamPP is switchable and can be turned on and off by using light of different wavelengths.

**Figure 5.**
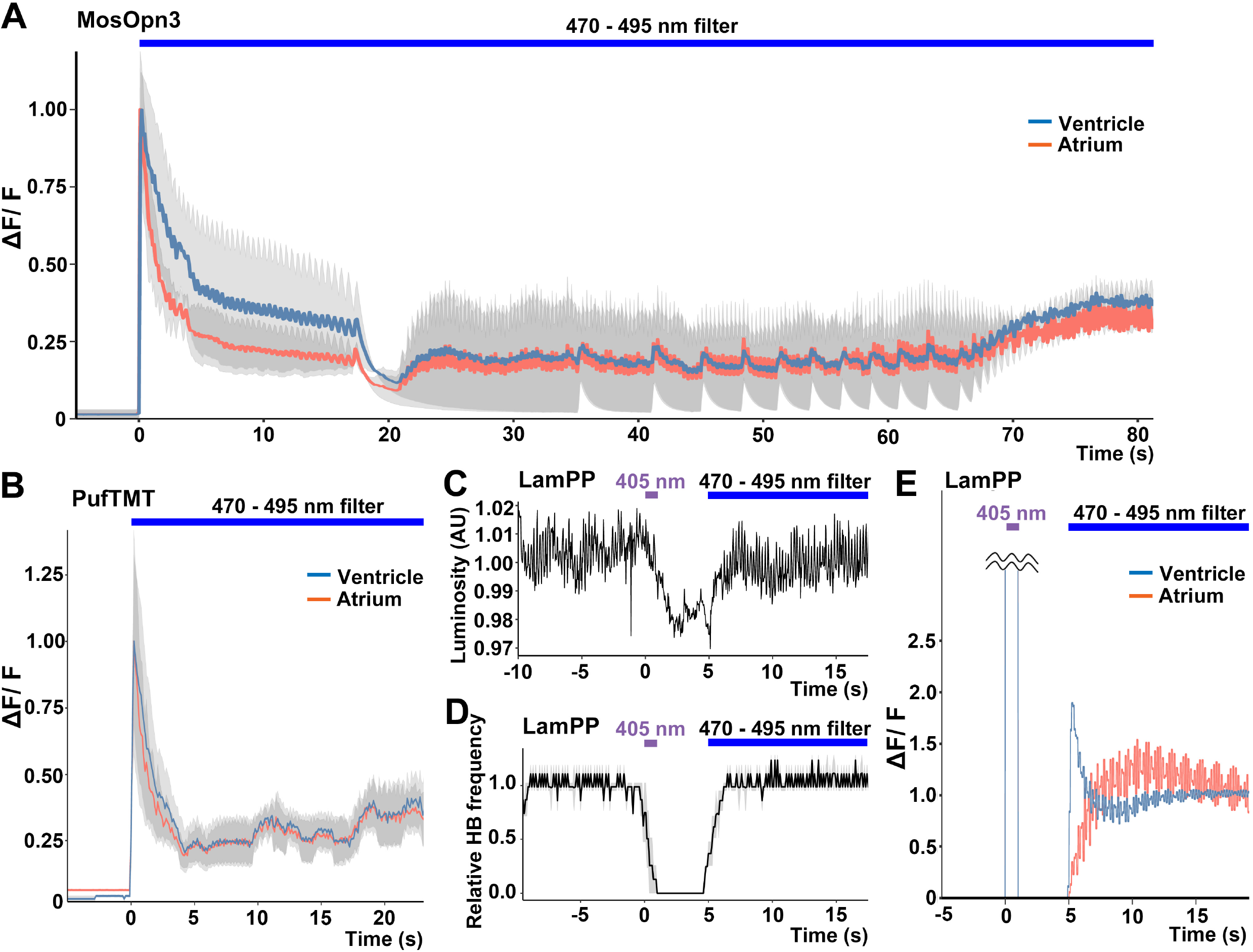
Switchable control of heartbeats by Gi/o-coupled bistable rhodopsins (A, B) Average changes in fluorescence of GCaMP6s (ΔF/F) of 4-dpf larvae expressing MosOpn3 (A) or PufTMT (B), and GCaMP6s in cardiomyocytes. The heart area was irradiated with a fluorescence detection filter (excitation 470-495 nm, emission 510-550 nm) for the indicated period (*n* = 2 for MosOpn3, *n* = 4 for PufTMT). ΔF/F was calculated as a ratio to the fluorescence intensity of GCaMP6s at the start of stimulation. (C, D) HB monitoring by luminosity (AU) change (C) and average of relative HB frequency (*n* = 2) (D) of 4-dpf larvae expressing LamPP in cardiomyocytes. The heart area was irradiated with 405 nm light (0.5 mW/mm^2^) for 1 s and then with a fluorescence detection filter (470-495 nm light) at the indicated period. (E) Changes in ΔF/F of GCaMP6s of a larva expressing LamPP and GCaMP6s in the heart. The heart area was irradiated with 405 nm light (0.5 mW/mm^2^) for 1 s and then with a fluorescence detection filter (470-495 nm light) at the indicated period. ΔF/F was calculated as the ratio to the fluorescence intensity of GCaMP6s at the steady state (after the resumption of HBs). Blue and red lines indicate ΔF/F in the ventricle and atrium (A, B, E).

### Gi/o-coupled rhodopsins suppress the heart’s function through an inward-rectifier K^+^ channel

To examine whether the optogenetic activity of MosOpn3, PufTMT, and LamPP depends on the activation of a Gi/o-type G protein, we treated the Tg fish expressing these rhodopsins with pertussis toxin (PTX), which induces ADP-ribosylation of Gαi and inhibits Gαi activity. We compared cardiac arrest time between PTX-treated fish and nontreated fish. Light-dependent activation of MosOpn3, PufTMT, or LamPP induced cardiac arrest, although the effect tended to weaken as the number of irradiation trials increased. This reduction in the effect of light stimulation might be due to residual activity of Gi/o-mediated signaling after repeated light stimulation trials. The cardiac arrest effect of these Gi/o-coupled rhodopsins was strongly suppressed by PTX treatment (Figure 6A, B, C, D, Movie 11), suggesting that optogenetic activity of these Gi/o-coupled rhodopsins requires the activation of the Gαi/o subunit.

**Figure 6.**
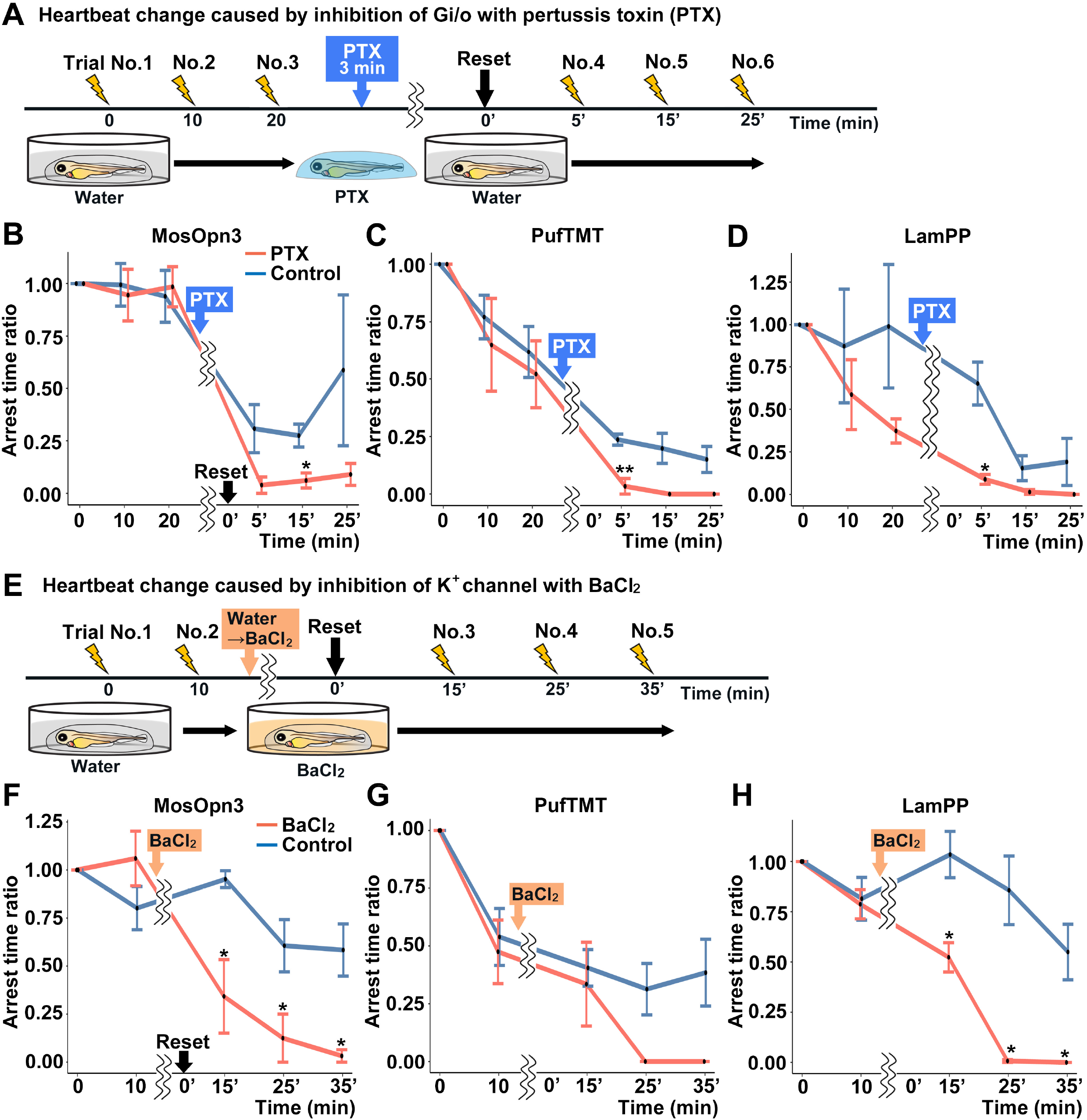
Gi/o and inward-rectifier K^+^ channel-dependent cardiac arrest by Gi/o-coupled bistable rhodopsins. (A) Time course of light irradiation and treatment with pertussis toxin (PTX) (min, minutes). 4-dpf Tg larvae expressing MosOpn3, PufTMT, or LamPP in cardiomyocytes were used. After three trials of light stimulation of the heart area in larvae embedded in agarose, the larvae were treated with PTX for 3 min and embedded in agarose again and subjected to three subsequent light stimulation trials. In each trial, the heart area was irradiated with appropriate wavelengths of light at an intensity of 0.5 mW/mm^2^ for 1 s, and cardiac arrest time was measured. The ratio to cardiac arrest time during the first trial was calculated (arrest time ratio). Light of wavelengths 520, 470, and 405 nm was used to stimulate MosOpn3, PufTMT, and LamPP, respectively. (B, C, D) Effect of PTX treatment on cardiac arrest induced by MosOpn3 (B), PufTMT (C), and LamPP (D). Average arrest time ratio of larvae expressing MosOpn3 (B), PufTMT (C), or LamPP (D) is shown in graphs. Larvae that were not treated with PTX were used as controls. Four treated and four non-treated control larvae were analyzed for each opto-tool. * *p* < 0.05, \*\**p* < 0.01, Welch’s *t* test. (E) Time course of light irradiation and treatment with BaCl_2_. After two trials of light stimulation of the heart area in larvae embedded in agarose, the larvae were treated with BaCl_2_ (or with water) and subjected to three subsequent light stimulation trials. In each trial, the heart area was irradiated with light of appropriate wavelengths at an intensity of 0.5 mW/mm^2^ for 1 s. Cardiac arrest time was measured and the arrest time ratio was calculated. (F, G, H) Effect of BaCl_2_ treatment on cardiac arrest induced by MosOpn3 (F), PufTMT (G), and LamPP (H). Average arrest time ratio of larvae expressing MosOpn3 (F), PufTMT (G), or LamPP (H) is shown in graphs. Larvae that were not treated with BaCl_2_ were used as controls. Four treated and four nontreated control larvae were analyzed for each opto-tool. \**p* < 0.05, Welch’s *t* test.

Gi/o-coupled GPCRs are known to suppress adenylyl cyclase (AC) and reduce intracellular cAMP. They are also known to hyperpolarize cells by increasing K^+^ efflux through an inward-rectifier K^+^ channel. To distinguish these two mechanisms, we treated Tg fish with BaCl_2_, an inhibitor of inward-rectifier K^+^ channels, and compared cardiac arrest time between incubation with BaCl_2_ and water (control). The light stimulusdependent cardiac arrest by MosOpn3, PufTMT, and LamPP was suppressed by incubation with BaCl_2_ (Figure 6E, F, G, H, Movie 12). The data suggest that the optogenetic activity of these Gi/o-coupled rhodopsins in the heart is dependent on the inward-rectifier K^+^ channel.

## Discussion

### Availability of animal bistable rhodopsins

We examined the optogenetic activities of G protein-coupled bistable rhodopsins derived from various vertebrate and invertebrate animals in zebrafish neurons and cardiomyocytes. We found that Gq-coupled SpiRh1 and its derivative SpiRh[S186F] could activate Gq-mediated signaling in reticulospinal V2a neurons. Gi/o-coupled MosOpn3, PufTMT, and LamPP could inhibit heart function when stimulated by light stimulation. Given that these bistable rhodopsins are sensitive to stimulating light of diverse wavelengths, they may be useful for manipulating various cell and tissue functions *in vivo* using light of different wavelengths. Animal bistable rhodopsins are endogenously expressed in various regions of the brain including photoreceptive tissues such as pineal and parapineal organs (*Kawano-Yamashita et al., 2011, Kawano-Yamashita et al., 2020, Kawano-Yamashita et al., 2015, Kawano-Yamashita et al., 2007, Koyanagi et al., 2004, Koyanagi et al., 2015, Shen et al., 2021, Wada et al., 2012, Wada et al., 2021, Wada et al., 2018*). If a wide area of the brain of Tg fish is irradiated with white light, it may also activate endogenous bistable rhodopsins in addition to transgene-expressed rhodopsins and affect the functions of neurons or other tissues. It is, therefore, important to compare the effects of light stimulation between transgenic and non-transgenic control fish. In this study, patterned illumination of a specific area of the brain or heart with light of selected wavelength lights enabled us to control the functions of target cells in Tg but not in non-transgenic fish (Figure 3A, 4D).

The bistable rhodopsins used in this study were photosensitive and functional without the addition of retinal derivatives *in vivo*. The bistable rhodopsins that bind to 11-*cis* retinal convert into an active state having all-*trans* retinal upon light absorption, and revert to the original inactive state by subsequent light absorption (*Koyanagi et al., 2021, Koyanagi and Terakita, 2014, Terakita, 2005, Terakita et al., 2015*). This bleach-resistant property confers activity to these bistable rhodopsins in non-photoreceptor cells. MosOpn3 was reported to bind to 13-*cis* retinal (*Koyanagi et al., 2013*). The 13-*cis* retinal-binding property of MosOpn3 assisted it to function in extraocular tissues since 13-*cis* retinal is generated in thermal equilibrium with the all-*trans* form, so 13-*cis* retinal is ubiquitously present (*Terakita et al., 2015*). In any case, the bistable rhodopsins can be activated by light in various types of cells other than retinal cells.

### Light-dependent activation with Gq-coupled rhodopsins

We observed robust neuronal activation and an increase in Ca^2+^ in reticulospinal V2a neurons expressing Gq-coupled SpiRh1 and SpiRh1[S186F] (Figure 3). We also observed tail movements after light stimulation of the axons of reticulospinal V2a neurons in the spinal cord with SpiRh1 (data not shown), suggesting that SpiRh1 is localized in axons, soma, and dendrites, and mediates Gq-mediated signaling. PLCβ mediates Gq-coupled signaling and produces IP_3_ and DAG from PIP_2_, which subsequently induces the release of Ca^2+^ from the ER and activates PKC and calmodulin kinases (CaMKs). It has been reported that binding of acetylcholine to the Gq-coupled muscarinic receptor (M1) activates non-selective cation channels and inhibits M-type K^+^ channels, inducing depolarization for a long period (*Fisahn et al., 2002, Fraser and MacVicar, 1996, Hofmann and Frazier, 2010, McQuiston and Madison, 1999, Yue and Yaari, 2004*). The inhibition of M-type K^+^ channels is considered to involve the PLCβ-mediated reduction of PIP_2_ (*Brown, 2010*). The same mechanism might be involved in neural activation, i.e. depolarization and generating action potentials, by SpiRh1 and SpiRh1[S186F]. It is also plausible that when Ca^2+^ increased, activated PKC and CaMKs phosphorylate cation channels, including neurotransmitter receptors, and this may also contribute to neural depolarization. This depolarization further leads to activation of voltage-dependent calcium channels. Consistent with this event, a burst in Ca^2+^ was observed upon generation of action potentials after stimulation with SpiRh1 and SpiRh1[S186F] (Figure 3, Movie 4, 5). Although Gq-coupled PLCβ-mediated signaling takes a longer time than channelrhodopsin-mediated signaling to activate neurons, this feedforward mechanism likely contributes to robust and long-lasting neuronal activation.

Two types of rhodopsins, channelrhodopsin and Gq-coupled rhodopsins, were shown to activate reticulospinal V2a neurons. Whereas photoactivation of channelrhodopsins immediately induced depolarization following cation influx, photoactivation of Gq-coupled rhodopsins induced a delayed increase in Ca^2+^ and neuronal activation. Similar neural activation takes place by binding of neurotransmitters to their receptors. For example, binding of glutamate to ion channel-type AMPA receptors and GPCR-type metabotropic receptors (GRMs), which are often present on the same postsynaptic membrane, likely induces immediate depolarization and a delayed Ca^2+^/depolarization pathway. Given that the two signals have different roles in neural circuit function, SpiRh1 and SpiRh1[S186F], together with channelrhodopsins, may be helpful in distinguishing the roles of these two signals in neural circuit function.

Light stimulation with SpiRh1 in cardiomyocytes did not apparently affect heart function (Table 1). The contraction of heart muscles requires an increase in intracellular Ca^2+^. It remains elusive whether SpiRh1 activation does not induce a sufficient increase in Ca^2+^ to affect heart function, or whether cooperation of action potentials together with an increase in Ca^2+^ is required for optic control of heart function. Future studies with calcium and voltage imaging and/or optogenetic activation of multiple pathways may clarify this issue.

### Light-dependent activation with Gi/o-coupled rhodopsins

Light stimulation of Gi-coupled rhodopsins MosOpn3, PufTMT, and LamPP in the heart induced cardiac arrest (Figure 4). This activity was inhibited by treatment with PTX and BaCl_2_ (Figure 6), suggesting that the Gi/o-coupled rhodopsins suppress neuronal activity by K^+^ channel-mediated hyperpolarization, which is mediated by the Gβγ subunit. It was previously reported that MosOpn3 and LamPP decreased neuronal excitability by coupling to inward-rectifier K^+^ (GIRKs) channels, but some form of MosOpn3-mediated inhibition of neural transmission was GIRK-independent (*Mahn et al., 2021*). It is possible that the PTX and BaCl_2_ treatments might affect functional expression of endogenous Gi/o-coupled GPCRs and indirectly affect the activity of the Gi/o-coupled rhodopsins. However, considering the complete suppression of light-induced cardiac arrest (Figure 6), these Gi/o-coupled rhodopsins likely suppress the heart’s function through inward-rectifier K^+^ channels in cardiomyocytes. As Gi/o-coupled GPCRs also regulate intracellular cAMP level via AC regulation, light stimulation of MosOpn3, PufTMT, LamPP, or ZPP1 reduced cAMP levels in HEK293S cells (Figure 1). The Gi/o -mediated control of cell functions may depend on cell type and subcellular location. We expressed MosOpn3, PufTMT, and LamPP in reticulospinal V2a neurons, but light activation of these Gi-coupled rhodopsins did not suppress spontaneous tail movements (Table 1). The inability to suppress tail movements may be due to slow activation of Gi/Go-mediated signaling by these bistable rhodopsins or the lack of other components in V2a neurons. Optimization of these tools and stimulation methods may be necessary, depending on cell type.

### Bistable nature of G-coupled rhodopsins

A short duration of light stimulation (1 s) of the heart expressing MosOpn3 or PufTMT induced cardiac arrest, resumed HBs after 10 s, and returned to a steady state after a few minutes (Figure 4), while prolonged light irradiation returned HBs to a steady state in a shorter time after cardiac arrest than short light irradiation (Figure 5). As the wavelengths of light effective for activation and inactivation were close for MosOpn3 and PufTMT, light irradiation likely induced both activation and inhibition of these Gi-coupled bistable rhodopsins. In contrast, the light wavelengths for activation and inactivation were apart for LamPP, which is switchable between these two states (*Copits et al., 2021, Koyanagi et al., 2004, Rodgers et al., 2021*). Consistent with this, cardiac arrest was induced by 405 nm light with LamPP, while irradiation of around 470 nm light resumed HBs (Figure 5C, D, E), suggesting that LamPP can be turned on and off by different wavelengths of light in the heart. Like LamPP, ZPP1 has different light wavelengths for activation and inactivation (Table 1). However, photoactivation of ZPP1 resulted in only a short period of cardiac arrest and its photosensitivity did not recover for a few hours. The photoproduct (active form) of ZPP1 might not be stable (i.e., it might release the chromophore easily) compared to that of MosOpn3, PufTMT, and LamPP in zebrafish cardiomyocytes.

Since Gq-coupled SpiRh1, and Gi-coupled MosOpn3, PufTMT, and LamPP are bistable rhodopsins, their photoproducts, which activate G protein-mediated signaling, are considered to be stable unless they receive inactivating light. The tail movements stopped several seconds after stimulation with SpiRh1 and SpiRh1[S186F], and HBs resumed a few minutes after stimulation with MosOpn3, PufTMT, and LamPP (Figure 2–4), suggesting that activity of the bistable rhodopsins gradually reduced after transient stimulation. Therefore, there are likely to be mechanisms that inactivate bistable rhodopsins other than the photo-dependent reversal from an active to an inactive form. They may not involve the release of all-*trans* retinal, but instead involve the phosphorylation-dependent binding of β-arrestin to rhodopsins and the β-arrestin-mediated internalization of rhodopsins (*Kawano-Yamashita et al., 2011*). In any case, by using G protein-coupled bistable rhodopsins with different properties (activating/inactivating light wavelengths, stability, etc.), the functions of cells and tissues can be finely controlled by light stimulation.

### Utility of bistable rhodopsin to study cell and tissue functions

Optogenetic tools that are proven to be useful in mammals are also effective in zebrafish, and *vice versa*. The bistable rhodopsin tools that we designed are effective in zebrafish, but are also active in HEK293S cells (Table 1). It is most likely that when they are expressed in mammalian tissues, they can be used to optogenetically manipulate GPCR signaling *in vivo*. In this study, the expression plasmids for bistable rhodopsins were constructed to express tagged rhodopsin and P2A-TagCFP by the Gal4-UAS system in specific types of zebrafish cells. The expression plasmids can be easily modified for other species. As small epitope-tagged bistable rhodopsins were more active than fluorescent protein-fused rhodopsins (Figure 1), they could also be more active in cells of other species, including mammals.

The genome of a single vertebrate species contains hundreds of GPCR genes. Many GPCRs function as receptors for sensations (e.g. odorant and taste receptors), and some function as receptors of some endogenous ligands (*Pierce et al., 2002*). There are also many GPCR signals whose role *in vivo* is not yet known. In the nervous system, GPCRs function as metabotropic receptors for neurotransmitters and neuromodulators, and are involved in neuronal functions such as synaptic plasticity, involving long-term potentiation (LTP) or depression (LTD) in neural circuits (*Reiner and Levitz, 2018*). Optogenetic manipulation of individual GPCR signaling should lead to a better understanding of their roles in synaptic plasticity and neural circuits. GPCRs also play important roles in regulating the function of internal organs (*de Lucia et al., 2018, Pierce et al., 2002, Rockman et al., 2002*). The G-coupled bistable rhodopsins analyzed in this study may be useful tools for the optogenetic control of various cell and tissue functions.

## Material and methods

### Key resources table

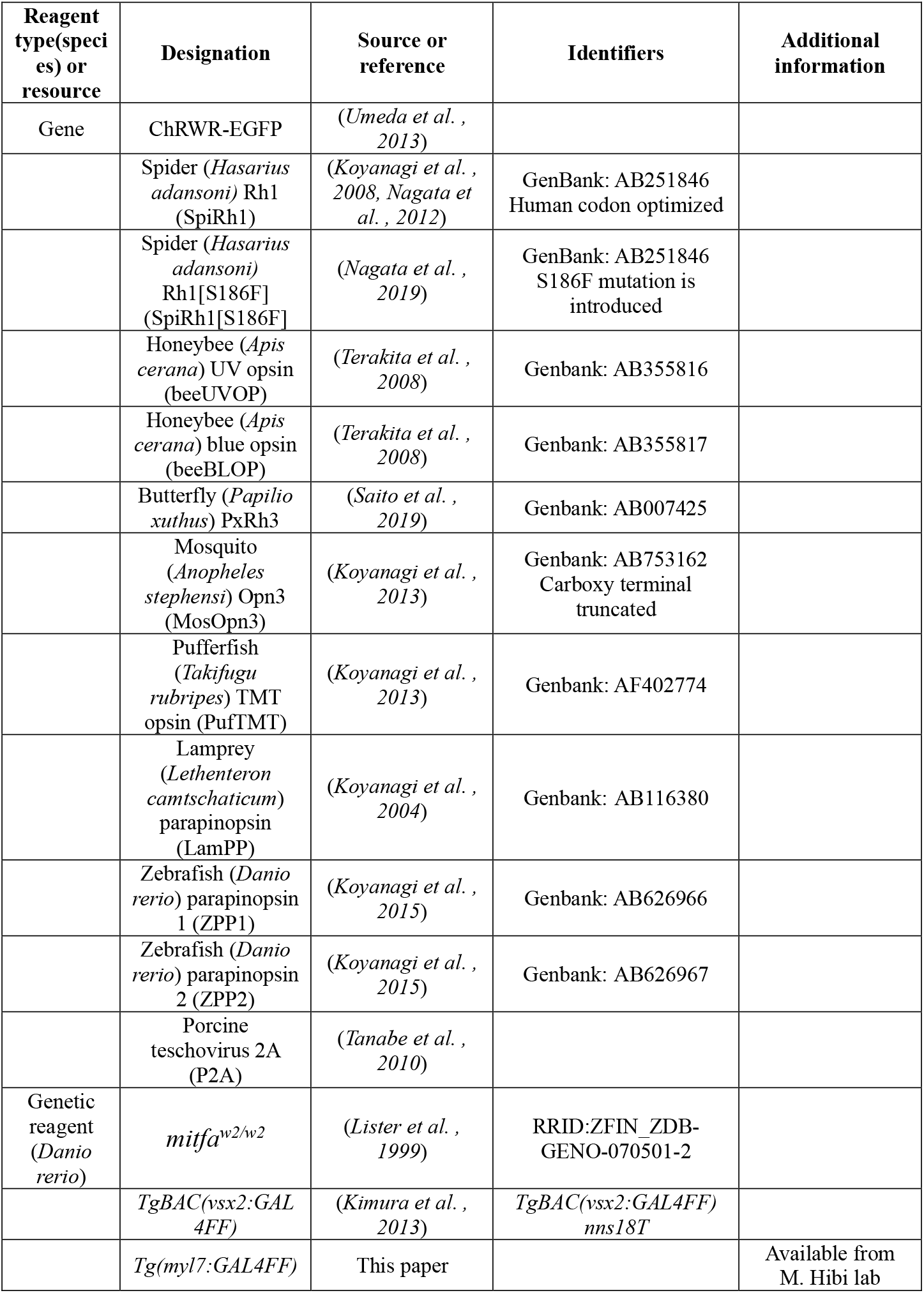
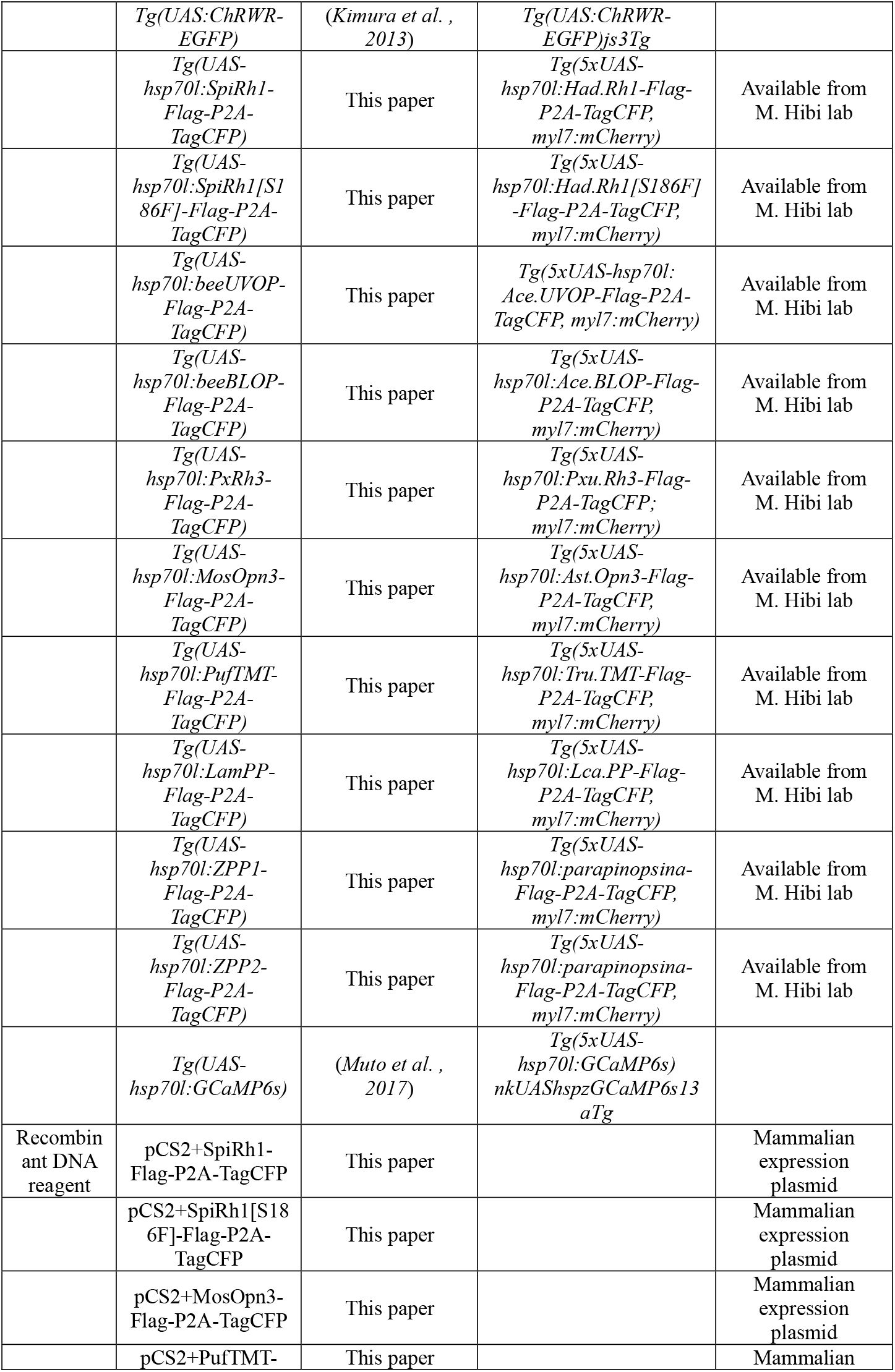
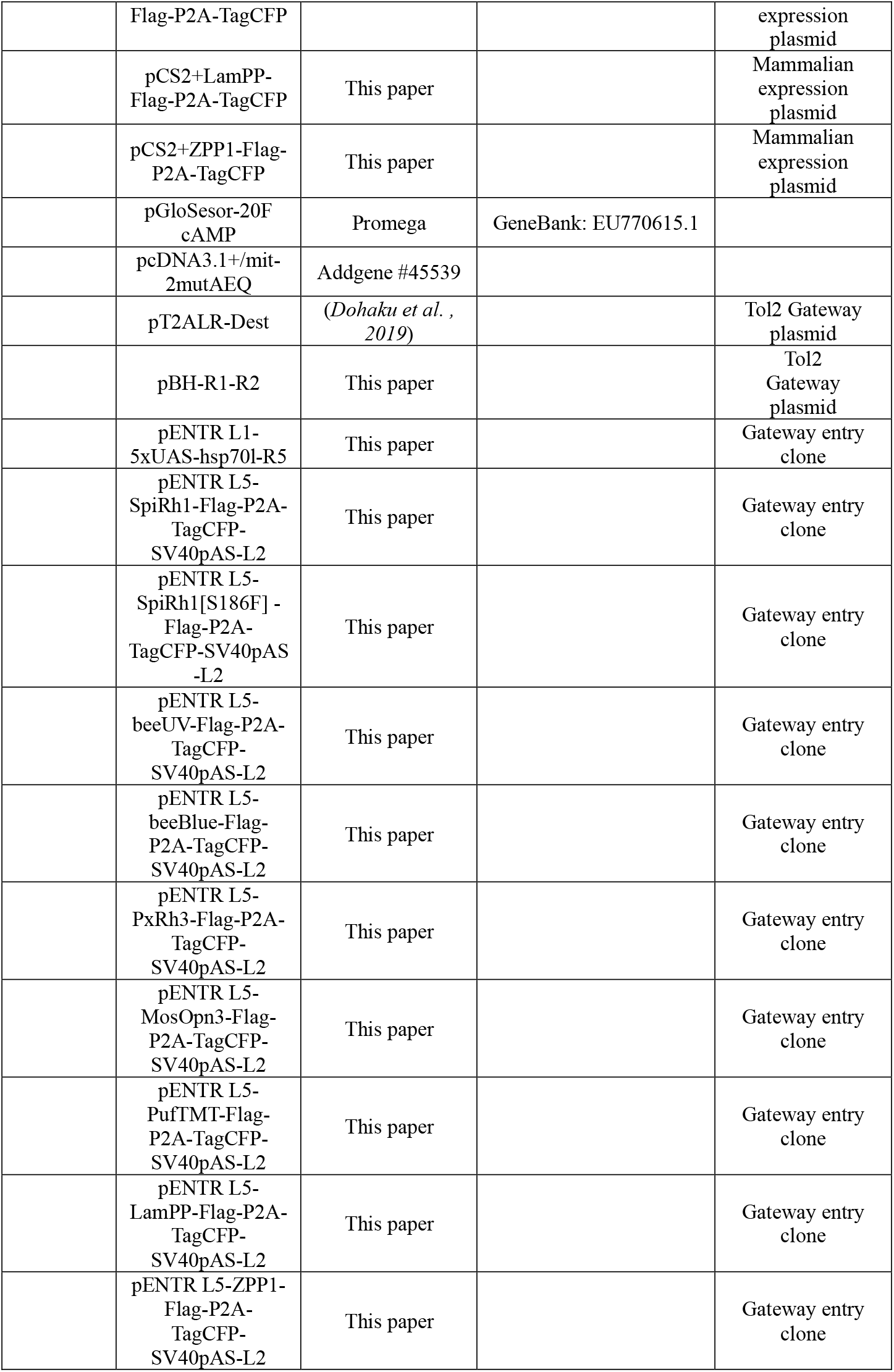
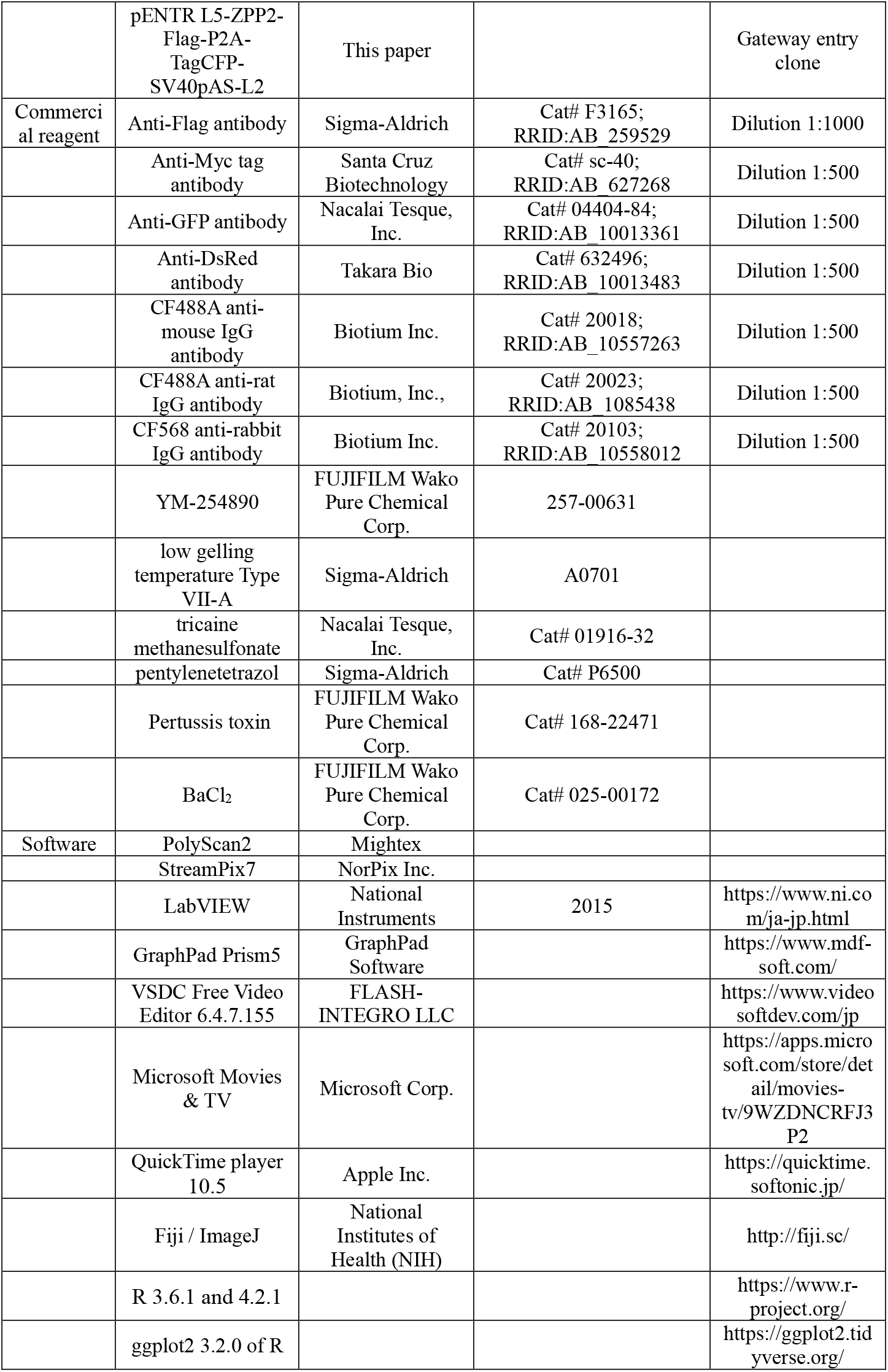
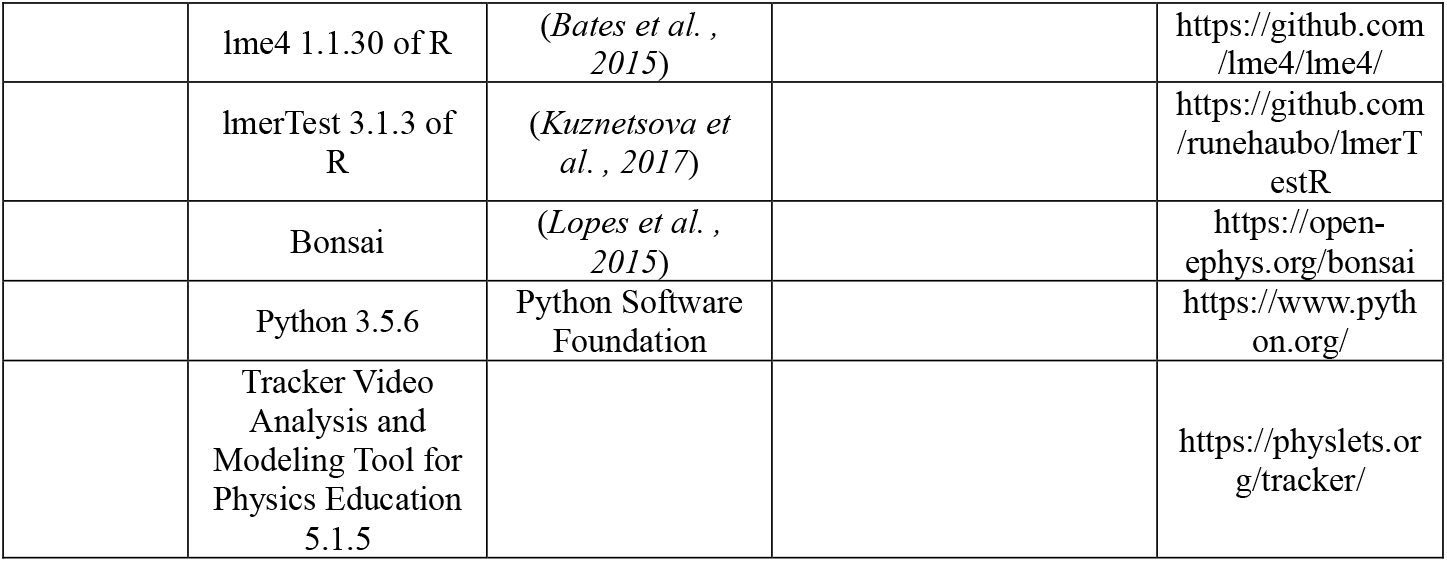

### Ethics statement

The animal experiments in this study were approved by the Nagoya University Animal Experiment Committee and were conducted in accordance with the Regulation on Animal Experiments from Nagoya University.

### Bioluminescent reporter assays for Ca^2+^ and cAMP

The intracellular cAMP and Ca^2+^ levels in rhodopsin-expressing HEK293S cells were measured using the GloSensor cAMP assay and the aequorin assay, respectively, as described previously (*Bailes and Lucas, 2013*). The pGloSensor-20F cAMP plasmid (Promega) was used for the GloSensor cAMP assay. The wild type aequorin obtained by introducing two reverse mutations into the plasmid [pcDNA3.1+/mit-2mutAEQ] (Addgene #45539) (*de la Fuente et al., 2012*) was used for the aequorin assay. The rhodopsin expression plasmids were constructed based on pCS2+ (see the section of Zebrafish) and used for transfection. For Gαq inhibition, YM-254890 (FUJIFILM Wako Pure Chemical Corp., 257-00631, Osaka, Japan) was added (1 μM) 5 min before the measurement. Green (500 nm) and violet (410 nm) LED lights were applied for 5 s in the GloSensor cAMP assay and for 1 s in the aequorin assay as light stimuli. Dual Head LED Light 505 nm (GB Life Science) and SPL-25-CC (REVOX, Inc.) were used for green and violet LED light stimulation, respectively.

### Zebrafish

All transgenic zebrafish lines in this study were generated using the *mitfa^w2/w2^* mutant (also known as *nacre*) line, which lacks melanophores (*Lister et al., 1999*). To generate plasmids for transgenesis expressing optogenetic tools, the open reading frame (ORF) of jumping spider (*Hasarius adansoni*) Rh1 (SpiRh1) (*Koyanagi et al., 2008, Nagata et al., 2012*), SpiRh1 S186F (*Nagata et al., 2019*), mosquito (*Anopheles stephensi*) Opn3 (*Koyanagi et al., 2013*), pufferfish (*Takifugu rubripes*) TMT opsin (*Koyanagi et al., 2013*), lamprey (*Lethenteron camtschaticum*) parapinopsin (*Koyanagi et al., 2004*), zebrafish (*Darnio rerio*) parapinopsin 1 and 2 (*parapinopsina* and *parapinopsinb* in ZFIN: https://zfin.org) (*Koyanagi et al., 2015*), honeybee (*Apis cerana*) UV and blue opsins (*Terakita et al., 2008*) or butterfly (*Papilio Xuthus*) PxRh3 (*Saito et al., 2019*) were amplified by PCR and subcloned to pCS2+ (pCS2+opto-tool) containing a Flag tag sequence, a 2A peptide sequence (P2A) from porcine teschovirus (PTV-1) (*Provost et al., 2007, Tanabe et al., 2010*), and TagCFP (Everon). pENTR L1-R5 entry vectors containing five repeats of the upstream activation sequence (UAS) and the *hsp70l* promoter (*Muto et al., 2017*), and pENTR L5-L2 vectors containing the ORF of the optogenetic tools and the polyadenylation site of SV40 (SV40pAS) from pCS2+opto-tools were generated by the BP reaction of the Gateway system. The UAS-hsp70l promoter (*Muto et al., 2017*) and optogenetic tool expression cassettes were subcloned to the Tol2 donor vector pBleeding Heart (pBH)-R1-R2 (*Dohaku et al., 2019*), which was modified from pBH-R4-R2 and contains mCherry cDNA and SV40 pAS under the *myosin, light chain 7, regulatory (myl7)* promoter (*van Ham et al., 2010*) by the LR reaction of the Gateway system. To make the Tol2 vector to express GAL4FF (a modified form of the yeast transcription factor GAL4) in the heart, an about 900-bp fragment of the promoter and a 5’ untranslated region (UTR) of *myl7* (from pKHR7) (*Hoshijima et al., 2016*), GAL4FF cDNA (*Asakawa et al., 2008*), and SV40pAS were subcloned to a Tol2 vector pT2ALR-Dest (*Dohaku et al., 2019*) by the Gateway system. To make Tg fish, 25 pg of the Tol2 plasmids and 25 pg of transposase-capped and polyadenylated RNA were injected into one-cell-stage embryos. The *Tg(UAS:opto-tool)* fish that expressed the optogenetic tools in a GAL4-dependent manner were crossed with *TgBAC(vsx2:GAL4FF);Tg(UAS:RFP) (Kimura et al., 2013), Tg(myl7:GAL4FF)*, or *Tg(elavl3:GAL4-VP16) (Kimura et al., 2013*) to express the tools in hindbrain reticulospinal V2a neurons, cardiomyocytes, and all postmitotic neurons, respectively. *Tg(UAS:ChRWR-EGFP)* was used as a positive control (*Kimura et al., 2013*). For Ca^2+^ imaging, *Tg(5xUAS-hsp70l:GCaMP6s) (Muto et al., 2017*) was used. Adult zebrafish were raised at 28.5°C with a 14 h light and 10 h dark cycle. Individual larvae used for behavioral experiments were kept in the dark except for the observation of fluorescence and light exposure experiments.

### Immunostaining

For immunostaining, anti-Flag antibody (1:500, mouse, Sigma-Aldrich, St. Louis, MO, USA, Cat# F3165; RRID:AB_259529), anti-Myc tag (MT) antibody (1:500, mouse, Santa Cruz Biotechnology, Dallas, TX, USA, Cat# sc-40; RRID:AB_627268), anti-GFP (1:500, rat, Nacalai Tesque, Inc., Kyoto, Japan, Cat# 04404-84; RRID:AB_10013361), and anti-DsRed (1:500, rabbit, Takara Bio, Shiga, Japan, Cat# 632496; RRID:AB_10013483) antibodies were used as primary antibodies. CF488A anti-mouse IgG (1:500, H + L, Biotium, Inc., Fremont, CA, USA, Cat# 20018; RRID:AB_10557263), CF488A anti-rat IgG (1:500, H + L, Biotium, Inc., Cat# 20023; RRID:AB_10854384), and CF568 anti-rabbit IgG (1:500, H + L, Biotium, Inc., Cat# 20103; RRID:AB_10558012) antibodies were used as secondary antibodies. Individual fish were placed in 1.5 mL Eppendorf tubes and fixed in 4% paraformaldehyde in PBS at 4°C for 1 h. The fixed samples were washed three times with PBST, treated with acetone for 12 min at room temperature, washed again three times with PBST and twice with PBS-DT. The solution was replaced with 5% goat serum in PBS-DT and was kept at room temperature for 1 h for blocking. Primary antibody was added to 5% goat serum in PBS-DT to achieve the dilution factor described above and incubated overnight at 4°C. The samples were washed with PBS-DT six times for 15 min each. The incubation in secondary antibody solution, 5% goat serum in PBS-DT with the above-mentioned dilution factor, was performed overnight at 4°C in the dark. After six washes of 15 min each in PBS-DT, the larvae were embedded in 1.5% agarose (low gelling temperature Type VII-A A0701, Sigma-Aldrich). Images were acquired using a confocal laser inverted microscope LSM700 (Carl Zeiss, Oberkochen, Germany). When taking images, the laser intensity was not changed by more than a factor of 2.

### Locomotion assay

3-dpf Tg larvae were quickly anesthetized with about 0.04% tricaine methanesulfonate (Nacalai Tesque, Inc., Kyoto, Japan, Cat# 01916-32) and embedded in 2.5% agarose in 1/10 Evans solution (134 mM NaCl, 2.9 mM KCl, 2.1 mM CaCl_2_, 1.2 mM MgCl_2_, and 10 mM Hepes; pH 7.8). The tail was set free by cutting the agarose around it. The agarose containing the embedded individual fish was placed in a 90-mm Petri dish filled with rearing water and kept under the microscope for 20 min to recover from anesthesia and to get used to the experimental environment which was followed by the first light exposure. For light stimulation, a patterned LED illuminator system LEOPARD (OPTOLINE, Inc., Saitama, Japan) and the control software PolyScan2 (Mightex, Toronto, Canada) was used. LEDs with wavelengths of 405, 488, 520, and 630 nm, which are the closest values to the maximum absorption wavelength of each optogenetic tool, were used. The irradiation intensity was adjusted to 0.4 mW/mm^2^. The irradiation area was 0.3 mm × 0.34 mm in the hindbrain (Figure 2A). Tail movements were captured by an infrared CMOS camera (67 fps, GZL-C1L-41C6M-C, Point Grey, Canada) mounted under the stage and StreamPix7 software (NorPix, Inc., Montreal, Canada) and analyzed by Tracker Video Analysis and Modeling Tool for Physics Education version 5.1.5. The timing of tail movement capture and light irradiation of the reticulospinal V2a neurons was controlled by a USB DAQ device (USB-6008, National Instruments, Austin, TX, USA) and programming software (LabVIEW, 2015, National Instruments). The irradiation stimulation was repeated six times every 20 min for 1 s (for G protein-coupled rhodopsins) or 100 ms (for ChRWR) with a minimum of eight individuals for each strain. The start and end times of tail movements were measured visually by StreamPix7 after the end of each trial. Trials in which swimming behavior was induced within 8 s after light stimulation were defined as induced trials. The percentage of induced trials was defined as locomotion rate, excluding trials in which swimming behavior was elicited before light stimulation. The time from the start of light irradiation to the first tail movement was defined as latency, and the time from the start of the first tail movement to the end of that movement was defined as duration. The maximum distance the tail moved from the midline divided by the body length was defined as strength. To examine the tools’ activity in the inhibition of locomotion, 4-dpf Tg larvae were pretreated with 15 mM pentylenetetrazol (Sigma-Aldrich, Cat# P6500) and spontaneous tail movements were induced by white LED light (peak 640 nm; Kingbright Electronic Co., Ltd., Taipei Hsien, Taiwan) powered by a DC power supply (E3631A; Agilent Technologies, Santa Clara, CA, USA) for 5 s. After 500 ms from the onset of tail movement, hindbrain reticulospinal V2a was stimulated with the patterned LED illuminator. Trials in which swimming behavior stopped within 1 s after light stimulation were defined as locomotioninhibition trials. The percentage of locomotion-inhibition trials was calculated and indicated in Table 1. Graphs were created with GraphPad Prism5 software (GraphPad Software, San Diego, CA, USA). All movies were created with VSDC Free Video Editor software version 6.4.7.155 (FLASH-INTEGRO LLC, Moscow, Russia) and Microsoft Movies & TV (Microsoft Corp., Redmond, WA, USA).

### Heartbeat experiments

4-dpf Tg larvae were quickly anesthetized with about 0.2% tricaine methanesulfonate and embedded in 4% agarose in 1/10 Evans solution. Larvae embedded in agarose were placed in a 90-mm Petri dish filled with water and kept under a microscope for 20 min for recovery from anesthesia. Light stimulation was performed as described in the section of the locomotion assay. Irradiation intensity was adjusted to 0.5 mW/mm^2^. The area of irradiation was 0.17 mm × 0.25 mm, including the heart. The heart area in the Tg fish expressing MosOpn3, PufTMT, or LamPP was irradiated for 1 s with light wavelength of 520, 470, and 405 nm, respectively, which are the closest values to the maximum absorption wavelength of each optogenetic tool. The HBs of larvae were captured by an infrared CMOS camera (67 fps) and recorded with StreamPix7, as described above. The irradiation trial was repeated six times every 10 min for one fish and a total of four larvae were analyzed for each strain. The video recordings of HBs were observed using the QuickTime player version 10.5 (Apple Inc., Cupertino, CA, USA). After opening videos with Fiji/ImageJ (National Institutes of Health, Bethesda, MD, USA), the entire heart was manually set as the region of interest (ROI), the luminosity (AU: arbitrary units) data in the ROI was used to create graphs of HBs using ggplot2 version 3.2.0 of R. The relative frequency of HBs was calculated by Bonsai (*Lopes et al., 2015*) and Python version 3.5.6 (Python Software Foundation, Wilmington, DE, USA). Graphs of the average of relative HB frequency were created by ggplot2 in R. The latency to cardiac arrest and the time to first resumption of HB were also measured. Graphs were created with GraphPad Prism5 software. All movies were created with VSDC Free Video Editor software. Simple HB experiments were also performed using a light source equipped with an MZ16 FA microscope and CFP (excitation light: 426-446 nm), GFP (460-500 nm), YFP (490-510 nm), and DSR filters (530-560 nm, Leica, Wetzlar, Germany), as indicated in Table 1.

### Treatment with pertussis toxin (PTX) or BaCl_2_

For PTX treatment, after the irradiation trial was repeated three times, the larvae were removed from agarose then immersed in a solution containing PTX (0.2 μg/mL, FUJIFILM Wako Pure Chemical Corp., Cat# 168-22471) for 3 min. After PTX treatment, larvae were embedded in agarose and placed on a Petri dish filled with deionized water. After larvae were kept in the Petri dish for 5 min, the heart area was irradiated three times every 10 min for 1 s (Figure 6). For control experiments of the pertussis toxin treatment, larvae were immersed in water instead of PTX solution for 3 min. For the BaCl_2_ treatment, 4-dpf larvae were embedded in agarose and placed in a Petri dish filled with water. After the irradiation trial was repeated twice, the water in the Petri dish was replaced with 1 mM BaCl_2_ (FUJIFILM Wako Pure Chemical Corp., Cat# 025-00172) solution. After larvae were kept in this solution for 15 min, the heart area was irradiated three times every 10 min for 1 s (Figure 6). For control experiments of the BaCl_2_ treatment, larvae were kept in water instead of BaCl_2_. After opening videos with QuickTime Player, cardiac arrest time was measured. Cardiac arrest ratio was calculated as the ratio to cardiac arrest time in trial 1, and plotted as a graph using ggplot2 of R.

### Ca^2+^ live imaging

Tg larvae expressing GCaMP6s with or without the opto-tool in reticulospinal V2a neurons or cardiomyocytes were quickly anesthetized with 0.04% tricaine methanesulfonate and embedded in 4% agarose in 1/10 Evans solution. A 130 W light source (U-HGLGPS, Olympus, Tokyo, Japan) with a fluorescence detection filter (excitation 470-495 nm, emission 510-550 nm, U-MNIBA3, Olympus) was used to observe the fluorescence of GCaMP6s. The same filter set was used to stimulate SpiRh1, MosOpn3, PufTMT, and LamPP. For Tg larvae expressing SpiRh1[S186F] or LamPP, the reticulospinal V2 neurons or the heart area were irradiated with 405 nm for 1 s with the patterned LED illuminator system. A CCD camera (ORCA-R2, Hamamatsu Photonics, Shizuoka, Japan) attached to the microscope was used to capture the GCaMP6s fluorescence images at 9 fps. After image acquisition of V2a neurons, the high intensity region from the hindbrain to the spinal cord was set as the ROI using ImageJ, and fluorescence intensity was measured. The relative change in fluorescence intensity (ΔF/F) was calculated by dividing the fluorescence intensity at each time point by the fluorescence intensity at the start of light stimulation for SpiRh1 or before stimulation (base line) for SpiRh1[S186F]. Graphs were created with GraphPad Prism5 software. After image acquisition for cardiomyocytes, videos of the heart were opened with Fiji/ImageJ, ROIs for the ventricle and atrium were set, and luminosity data were acquired. ΔF/F was calculated by dividing the fluorescence intensity at each time point by fluorescence intensity at the start of light stimulation for MosOpn3 and PufTMT, or by fluorescence intensity at the steady state (after HB resumption) for LamPP.

### Statistical analysis

Data were analyzed using R software package (versions 3.6.1 and 4.2.1). Statistical tests were applied as indicated in figure legends. All data in the text and figures are expressed as the mean ± standard error. A linear mixed-effects model was applied using R package ‘lme4’ version 1.1.30 and ‘lmerTest’ version 3.1.3 (*Bates et al., 2015, Kuznetsova et al., 2017*). The *p*-values of this model were computed with a *t* test based on the degrees of freedom with Scatterthwaite’s method, including time and strain as the fixed effects, and individuals as the random effect.

## Acknowledgements

We thank Hiromu Yawo, Shin-ichi Higashijima, Koichi Kawakami, and the National Bioresource Project for providing transgenic fish, Tamiko Itoh for managing fish mating and care, and Ryosuke Takeuchi for helping us analyze heartbeat experiments. We also thank the members of the Terakita and Hibi laboratories for helpful discussion.

---

**Figure S1.**
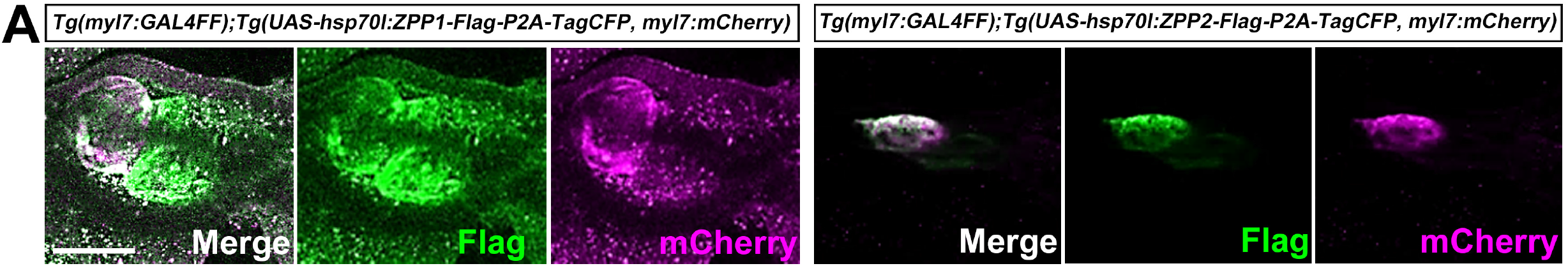
Expression of ZPP1 and ZPP2 in cardiomyocytes. Immunostaining of 4-dpf *Tg(myl7:GAL4FF);Tg(UAS:ZPP1-Flag-P2A-TagCFP, myl7:mCherry) Tg(myl7:GAL4FF); Tg(UAS:ZPP2-Flag-P2A-TagCFP, myl7:mCherry)* larvae, which were fixed and stained with anti-Flag (green) and anti-DsRed (mCherry: magenta) antibodies. Scale bar = 100 μm in (A).

## Movies

**Movie 1.** Tail movements in a larva expressing ChRWR-EGFP in reticulospinal V2a neurons.

The hindbrain in a 3-dpf *Tg(vsx2:GAL4FF);Tg(UAS-hsp70l:ChRWR-EGFP);Tg(UAS:RFP)* larva was stimulated with 470 nm light for 100 ms. The timing of light stimulation is indicated by a blue circle.

**Movie 2.** Tail movements in a larva expressing SpiRh1 in reticulospinal V2a neurons.

The hindbrain in a 3-dpf *Tg(vsx2:GAL4FF);Tg(UAS-hsp70l:SpiRh1-Flag-P2A-TagCFP, myl7:mCherry);Tg(UAS:RFP)* larva was stimulated with 520 nm light for 1 s. The timing of light stimulation is indicated by a green circle.

**Movie 3.** Tail movements in a larva expressing SpiRh1[S186F] in reticulospinal V2a neurons.

The hindbrain in a 3-dpf *Tg(vsx2:GAL4FF);Tg(UAS-hsp70l:SpiRh1-Flag-P2A-TagCFP; myl7:mCherry);Tg(UAS:RFP)* larva was stimulated with 405 nm light for 1 s. The timing of light stimulation is indicated by a purple circle.

**Movie 4.** Ca^2+^ imaging of hindbrain reticulospinal V2a neurons of a larva expressing SpiRh1 and GCaMP6s.

The hindbrain in 3-dpf *Tg(vsx2:GAL4FF);Tg(UAS-hsp70l:SpiRh1-Flag-P2A-TagCFP, myl7:mCherry);Tg(UAS-hsp70l:GCaMP6s)* was stimulated with a fluorescence detection filter (excitation 470-495 nm, emission 510-550 nm). The timing of stimulation is indicated by a blue square. GCaMP6s fluorescence was simultaneously monitored with the same filter.

**Movie 5.** Ca^2+^ imaging in hindbrain reticulospinal V2a neurons of a larva expressing SpiRh1[S186F] and GCaMP6s.

The hindbrain in 3-dpf *Tg(vsx2:GAL4FF);Tg(UAS-hsp70l:SpiRh1[S186F]-Flag-P2A-TagCFP, myl7:mCherry);Tg(UAS-hsp70l:GCaMP6s)* was stimulated with 405 nm light after detection of the baseline fluorescence. The timing of light stimulation is indicated by a purple circle. GCaMP6s fluorescence was monitored with a fluorescence detection filter (excitation 470-495 nm, emission 510-550 nm). The timings of the observations are indicated by blue squares.

**Movie 6.** Heartbeats in a larva expressing MosOpn3 in cardiomyocytes.

The heart area of *Tg(myl7:GAL4FF);Tg(UAS-hsp70l:MosOpn3-Flag-P2A-TagCFP, myl7:mCherry)* was stimulated with 520 nm light for 1 s. The timing of light stimulation is indicated by a green circle.

**Movie 7.** Heartbeats in a larva expressing PufTMT in cardiomyocytes.

The heart area of *Tg(myl7:GAL4FF);Tg(UAS-hsp70l:PufTMT-Flag-P2A-TagCFP, myl7:mCherry)* was stimulated with 470 nm light for 1 s. The timing of light stimulation is indicated by a blue circle.

**Movie 8.** Heartbeats in a larva expressing LamPP in cardiomyocytes.

The heart area of *Tg(myl7:GAL4FF);Tg(UAS-hsp70l:LamPP-Flag-P2A-TagCFP, myl7:mCherry)* was stimulated with 405 nm light for 1 s. The timing of light stimulation is indicated by a purple circle.

**Movie 9.** Ca^2+^ imaging in the heart of a larva expressing MosOpn3 and GCaMP6s.

The heart area of *Tg(myl7:GAL4FF);Tg(UAS-hsp70l:MosOpn3-Flag-P2A-TagCFP, myl7:mCherry);Tg(UAS-hsp70l:GCaMP6s)* was stimulated with a fluorescence detection filter (excitation 470-495 nm, emission 510-550 nm). GCaMP6s fluorescence was monitored with the same filter set. GCaMP6s fluorescence gradually decreased and became almost undetectable. At about 40 s, fluorescence recovered as the heart began to beat. A typical example is shown.

**Movie 10.** Heartbeat changes by stimulation with light of different wavelengths in a larva expressing LamPP in cardiomyocytes.

The heart area of *Tg(myl7:GAL4FF);Tg(UAS-hsp70l:LamPP-Flag-P2A-TagCFP, myl7:mCherry)* was stimulated with 405 nm light and a fluorescence detection filter (470-495 nm light). The timing of stimulation with 405 nm (1 s) and 470-495 nm light is indicated by a purple circle and a blue square, respectively. Cardiac arrest was induced by stimulation. Immediately after stimulation with 470-495 nm light, HBs were resumed. A typical example is shown.

**Movie 11.** Effect of pertussis toxin treatment on cardiac arrest induced by PufTMT activation.

4-dpf Tg larvae expressing PufTMT in cardiomyocytes were stimulated with 470 nm light for 1 s in the first three trials. Then they were treated with PTX (right panel) or remained untreated (left panel). Subsequently, they were stimulated with 470 nm light in trials 4-6. HBs in trials 1 and 4 are shown. The timing of stimulation is indicated by blue circles. Cardiac arrest was no longer observed after PTX treatment (PTX Tx; trial 4, right panel).

**Movie 12.** Effect on BaCl_2_ treatment on cardiac arrest induced by LamPP activation.

4-dpf Tg larvae expressing LamPP in cardiomyocytes were stimulated with 405 nm light in trials 1 and 2. They were then treated with BaCl_2_ (right panel) or remained untreated (left panel). Subsequently, they were stimulated with 405 nm light in trials 3-5. Heart movements in trials 1 and 5 are shown. The timing of stimulation is indicated by purple circles. Cardiac arrest was no longer observed after the BaCl_2_ treatment (BaCl_2_ Tx; trial 5, right panel).

